# A calibrated optogenetic toolbox of stable zebrafish opsin lines

**DOI:** 10.1101/2020.01.13.904185

**Authors:** P Antinucci, AS Dumitrescu, C Deleuze, HJ Morley, K Leung, T Hagley, F Kubo, H Baier, IH Bianco, C Wyart

## Abstract

Optogenetic actuators with diverse spectral tuning, ion selectivity and kinetics are constantly being engineered providing powerful tools for controlling neural activity with subcellular resolution and millisecond precision. Achieving reliable and interpretable *in vivo* optogenetic manipulations requires reproducible actuator expression and calibration of photocurrents in target neurons. Here, we developed nine transgenic zebrafish lines for stable opsin expression and calibrated their efficacy *in vivo*. We first used high-throughput behavioural assays to compare opsin ability to elicit or silence neural activity. Next, we performed *in vivo* whole-cell electrophysiological recordings to quantify the amplitude and kinetics of photocurrents and test opsin ability to precisely control spiking. We observed substantial variation in efficacy, associated with differences in both opsin expression level and photocurrent characteristics, and identified conditions for optimal use of the most efficient opsins. Overall, our calibrated optogenetic toolkit will facilitate the design of controlled optogenetic circuit manipulations.

## Introduction

Optogenetics has greatly advanced our ability to investigate how neural circuits process information and generate behaviour by allowing manipulation of neural activity with high spatio-temporal resolution in genetically-defined neurons (Miesenbock, 2009; Boyden, 2011; Miesenbock, 2011; Adamantidis *et al*., 2015; Boyden, 2015; Deisseroth, 2015; Deisseroth and Hegemann, 2017). The efficacy with which optogenetic actuators – such as microbial opsins – can control neuronal spiking *in vivo* depends on biophysical properties, expression level and membrane trafficking of the opsin, physiological properties of the target cell and the intensity profile of light delivered within scattering tissue.

Accordingly, two primary experimental requirements should be met to enable controlled and reproducible *in vivo* optogenetic circuit manipulations: (*i*) reproducible opsin expression levels (across cells and animals), with stable expression systems offering higher reliability and homogeneity than transient ones (Kikuta and Kawakami, 2009; Yizhar *et al*., 2011; Sjulson *et al*., 2016), and (*ii*) calibrated photocurrents recorded in target neurons (Huber *et al*., 2008; Li *et al*., 2019). While previous studies have compared the physiological effects of opsin activation in single cells using standardised conditions [e.g. (Berndt *et al*., 2011; Mattis *et al*., 2011; Prigge *et al*., 2012; Klapoetke *et al*., 2014; Berndt *et al*., 2016; Mardinly *et al*., 2018)], these comparisons were primarily performed *in vitro* or *ex vivo* using transient expression strategies.

In this study, we took advantage of the genetic accessibility and transparency of zebrafish (Arrenberg *et al*., 2009; Del Bene and Wyart, 2012; Arrenberg and Driever, 2013; Portugues *et al*., 2013; Forster *et al*., 2017) to generate nine stable transgenic lines for targeted opsin expression using the GAL4/UAS binary expression system (Scheer and Campos-Ortega, 1999; Asakawa and Kawakami, 2008) and quantitatively compare their efficacy for inducing or silencing neuronal spiking. We selected opsins that were reported to induce photocurrents with large amplitude [CoChR (Klapoetke *et al*., 2014), CheRiff (Hochbaum *et al*., 2014), ChR2(H134R) (Gradinaru *et al*., 2007), eArch3.0 (Mattis *et al*., 2011), GtACR1–2 (Govorunova *et al*., 2015)] and/or fast kinetics [Chronos, ChrimsonR (Klapoetke *et al*., 2014), eNpHR3.0 (Gradinaru *et al*., 2010)]. We first assessed the efficacy of these stable lines to control activity in intact neural populations via high-throughput behavioural assays at both embryonic and larval stages. Next, we made *in vivo* electrophysiological recordings from single low input-resistance motor neurons to calibrate photocurrents and test the ability of each line to elicit or silence spiking. We observed broad variation in behavioural response rates, photocurrent amplitudes and spike induction, likely due to differences in both opsin properties and expression levels. For the best opsin lines, we identified conditions that allowed control of individual action potentials within high-frequency spike trains. Overall, our toolkit will enable reliable and robust optogenetic interrogation of neural circuit function in zebrafish.

## Results

### Generation of stable transgenic lines for targeted opsin expression in zebrafish

To maximise the utility of our optogenetic toolkit, we used the GAL4/UAS binary expression system for targeted opsin expression in specific cell populations (Figure 1). We generated nine stable UAS lines for opsins having different ion selectivities and spectral tuning, fused to a fluorescent protein reporter (tdTomato or eYFP; Figure 1A and Supplementary File 1) (Asakawa *et al*., 2008; Arrenberg *et al*., 2009; Horstick *et al*., 2015). GAL4 lines were used to drive expression in defined neuronal populations, such as motor neurons (Figure 1B) (Scott *et al*., 2007; Wyart *et al*., 2009; Bohm *et al*., 2016). High levels of expression were achieved in most cases (Figure 1C), with only few opsins showing intracellular puncta suggestive of incomplete trafficking to the plasma membrane (CheRiff and GtACR2) or low expression (Chronos). To quantitatively compare opsin lines, we performed standardised behavioural tests at embryonic and larval stages (Figure 1D) and calibrated photocurrents and modulation of spiking in larval primary motor neurons (Figure 1E).

**Figure 1.**
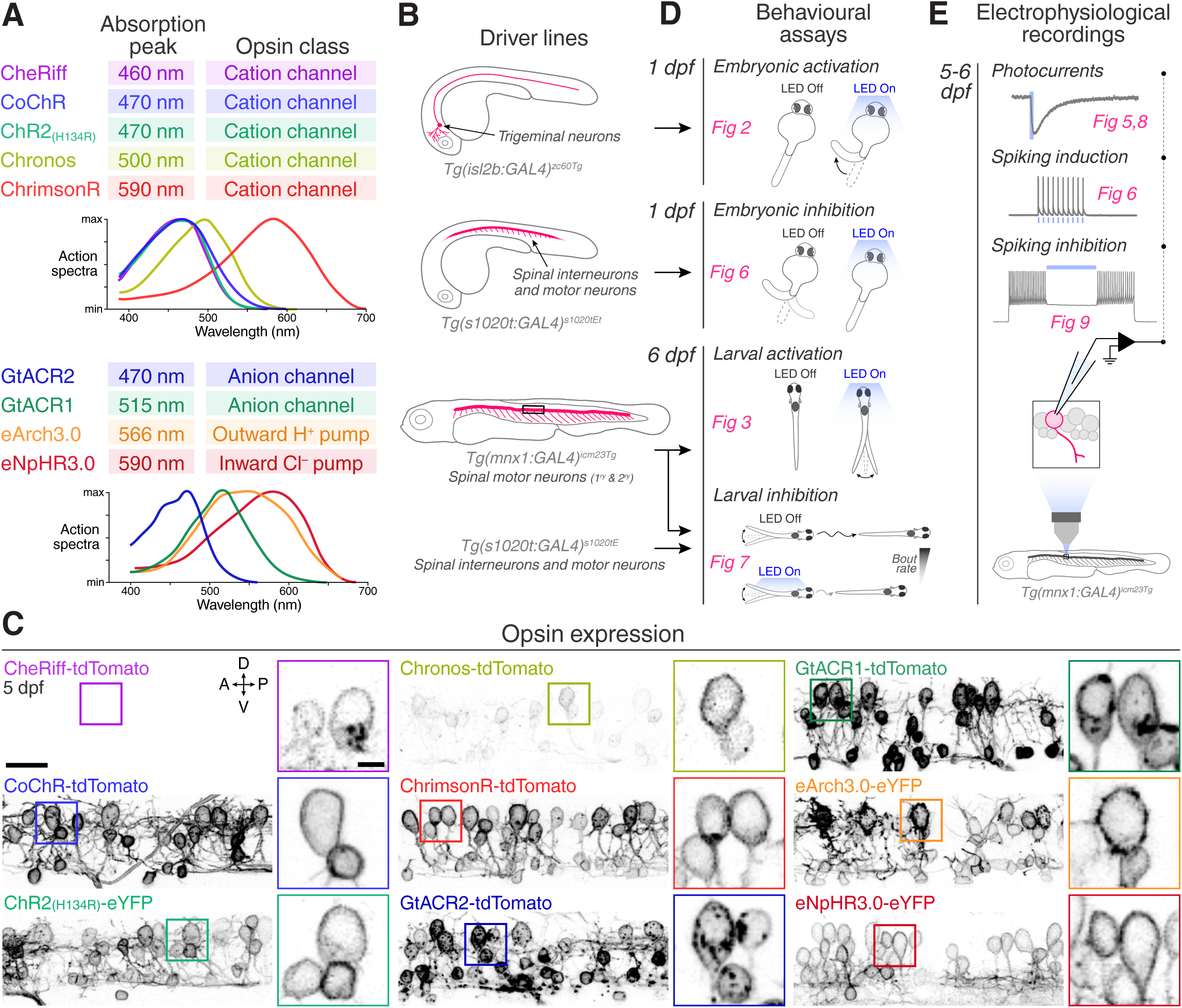
Toolkit for targeted opsin expression. **A** List of selected opsins, with spectral absorption and opsin class. **B** Schematics of expression patterns in the GAL4 transgenic driver lines used in this study. **C** Opsin expression in spinal neurons in *Tg(mnx1:GAL4;UAS:opsin-FP)* larvae at 5 dpf (for eNpHR3.0, the *s1020t:GAL4* transgene was used). Insets show magnified cell bodies to illustrate opsin membrane expression (for insets, brightness and contrast were adjusted independently for each opsin to aid visualisation). A, anterior; D, dorsal; P, posterior; V, ventral. Scale bar 20 *μ*m in large images, 5 *μ*m in insets. **D** Behavioural assays and corresponding figure numbers. **E** *In vivo* electrophysiological recordings and figure numbers.

### Escape behaviour triggered by optogenetic activation of embryonic trigeminal neurons

As a first test of our opsin lines, we evaluated their ability to activate embryonic neurons (Figure 2A– C), which are characterised by high input resistance (Drapeau *et al*., 1999; Saint-Amant and Drapeau, 2000). We used the *Tg(isl2b:GAL4)* transgene (Ben Fredj *et al*., 2010) to drive expression of opsins in the trigeminal ganglion (Figure 2B,C). In this class of somatosensory neuron, optogenetic induction of few spikes has been shown to reliably elicits escape responses (Douglass *et al*., 2008), characterised by high-amplitude bends of the trunk and tail (Kimmel *et al*., 1990; Saint-Amant and Drapeau, 1998; Sagasti *et al*., 2005). Brief pulses of light (5 or 40 ms-long) induced escape responses in embryos (28– 30 hours post fertilisation, hpf) expressing all cation- and anion-conducting channelrhodopsins (Figure 2C–E and Video 1), while no movement was elicited in opsin-negative siblings (Figure 2F,G and Figure 2–figure supplement 1,2; N = 69 ± 26 fish per group, mean ± SD). The excitatory effect of GtACRs suggests that increasing chloride conductance depolarises neurons at this developmental stage. For all opsins, response probability increased monotonically with light power (Figure 2F,G). Escape behaviour could also be evoked via transient opsin expression, in which animals were tested one day after injection of DNA constructs into single cell-stage *Tg(isl2b:GAL4)* embryos (Figure 2F). Some opsins showed higher response probability in transient transgenic animals (CheRiff, CoChR and GtACRs), likely due to higher expression levels.

**Figure 2.**
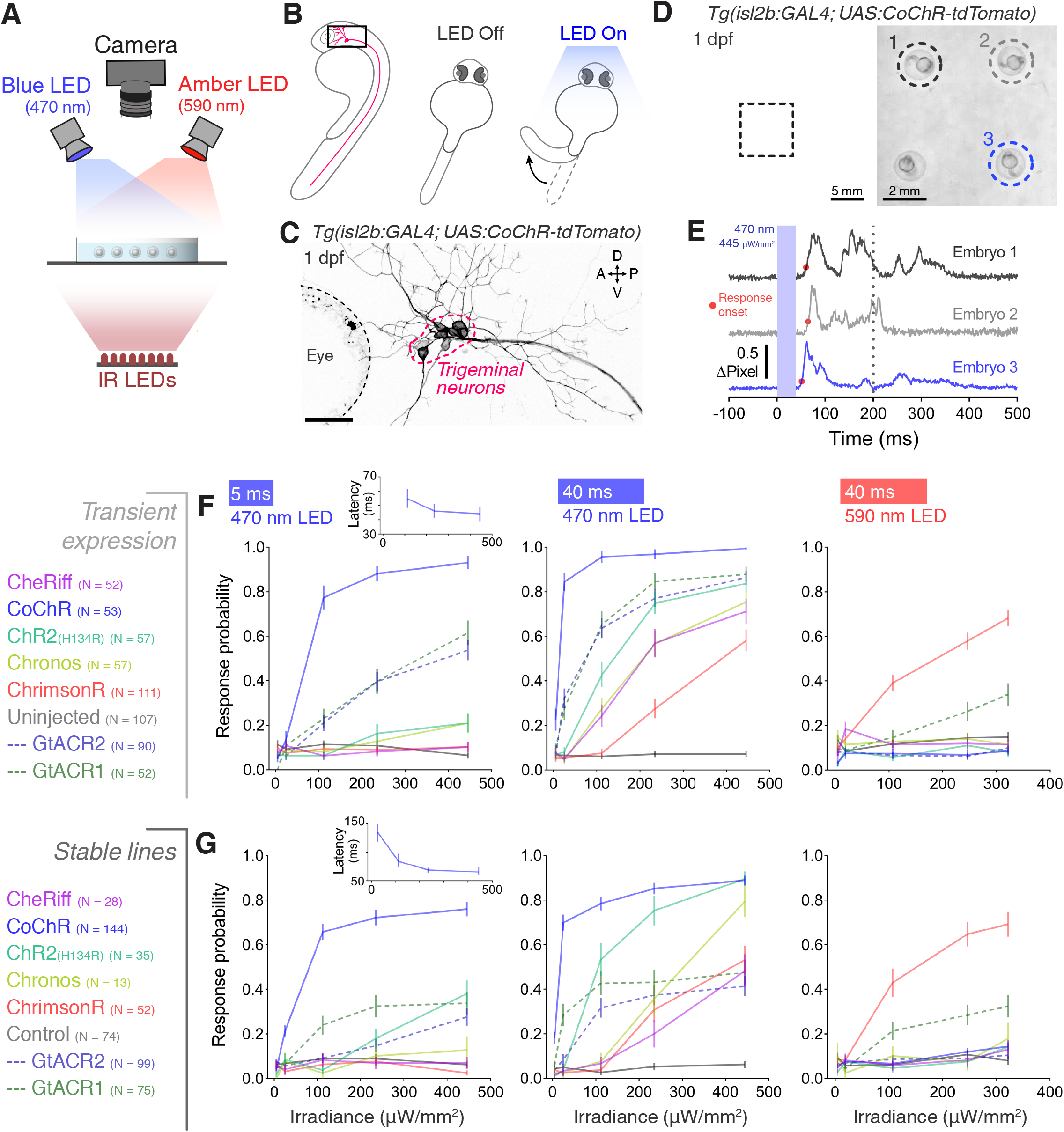
Optogenetic activation of embryonic trigeminal neurons triggers escape responses. **A** Experimental setup for optogenetic stimulation and behavioural monitoring. IR, infrared. **B** Schematic of behavioural assay. **C** Opsin expression in trigeminal neurons in a *Tg*(*isl2b:GAL4;UAS:CoChR-tdTomato)* embryo at 1 dpf. Imaging field of view corresponds to black box in (**B**). A, anterior; D, dorsal; P, posterior; V, ventral. Scale bar 50 μm. **D** *Tg(isl2b:GAL4;UAS:CoChR-tdTomato)* embryos positioned in individual agarose wells. Behaviour was monitored at 1,000 frames per second across multiple embryos (28–30 hpf; N = 69 ± 26 fish per opsin group, mean ± SD) subjected to 5 or 40 ms pulses of full-field illumination (470 or 590 nm, 4.5–445 *μ*W/mm^2^) with a 15 s inter-stimulus interval. **E** Optogenetically-triggered escape responses detected from ΔPixel traces in the 3 embryos indicated in (**D**). Dotted line indicates maximum latency (200 ms) for a response to be considered optogenetically-triggered. **F,G** Response probability for transient (**E**) or stable (**F**) transgenic embryos expressing different opsins (mean ± SEM, across fish). Insets show response latency for 5 ms blue light pulses in CoChR-expressing embryos (median ± 95% CI, across fish).

With blue light, CoChR elicited escapes at the highest response probability (65–100% at 112– 445 *μ*W/mm^2^; Figure 2F,G) and response latency decreased with increasing irradiance (insets in Figure 2F,G). As expected from its red-shifted absorption spectrum, ChrimsonR was the only cation channelrhodopsin to evoke escapes using amber light (∼70% response probability at 322 *μ*W/mm^2^; Figure 2F,G) (Klapoetke *et al*., 2014). Consistent with their respective red- and blue-shifted absorption spectra, GtACR1 triggered escapes upon amber and blue light stimulation whereas GtACR2 elicited responses only with blue light (Figure 2F,G) (Govorunova *et al*., 2015).

### Tail movements triggered by optogenetic activation of larval spinal motor neurons

Next, we compared the efficacy of cation channelrhodopsin lines to induce behaviour by activation of larval motoneurons, from which we would later record photocurrents. We used the *Tg(mnx1:GAL4)* transgene (Bohm *et al*., 2016) to target expression to spinal motor neurons (Figure 3A,B) and subjected head-restrained zebrafish (6 days post fertilisation, dpf; N = 28 ± 8 fish per group, mean ± SD) to either single light pulses (2 or 10 ms-long) or pulse trains at 20 or 40 Hz (Figure 3C,D and Video 2,3) while monitoring tail movements.

**Figure 3.**
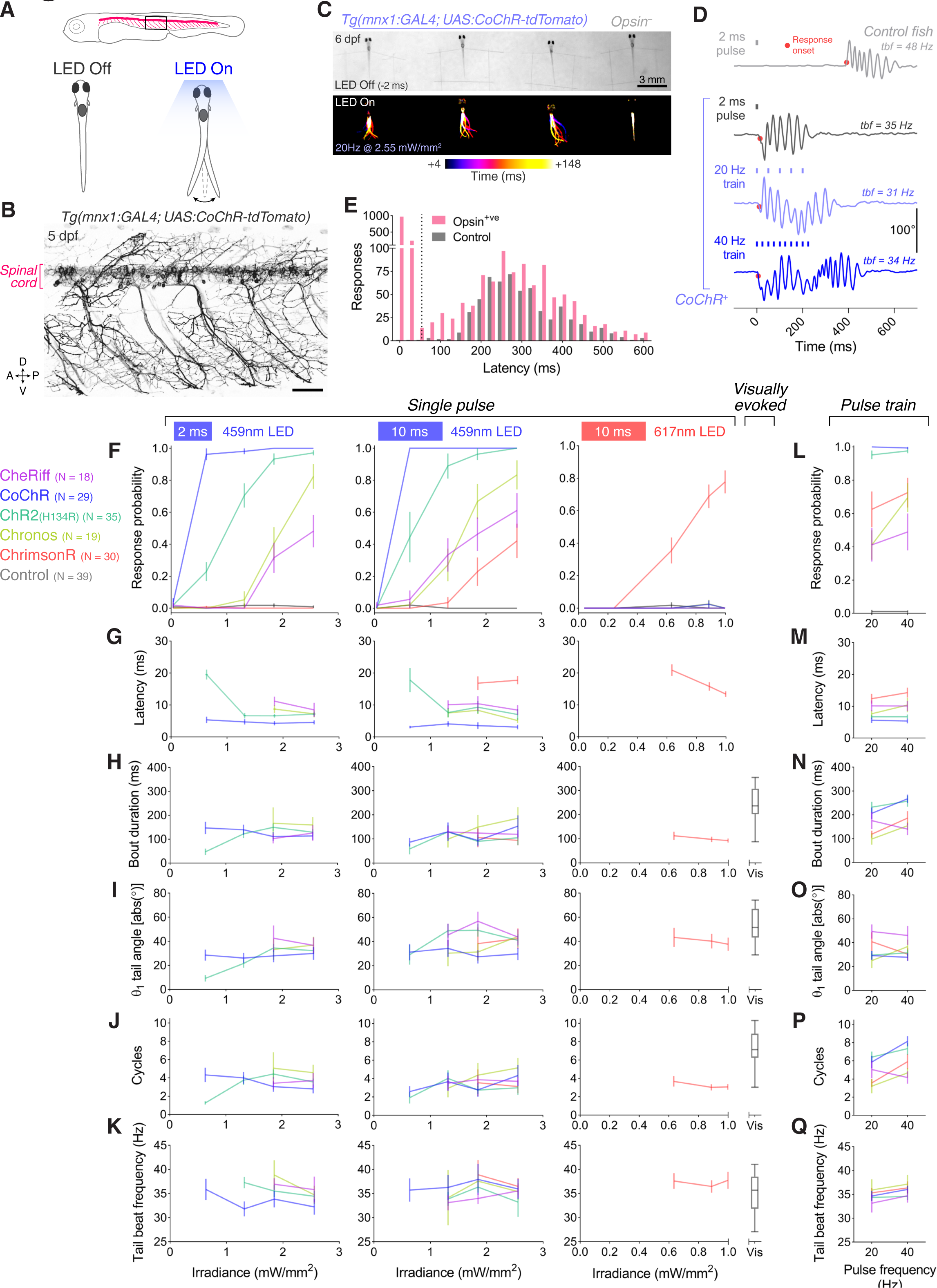
Optogenetic activation of larval spinal motor neurons triggers tail movements. **A** Schematics of behavioural assay. Head-restrained, tail-free larvae (6 dpf; N = 28 ± 8 fish per opsin group, mean ± SD) were exposed to 2 or 10 ms pulses of light (459 or 617 nm, 0.04–2.55 mW/mm^2^) with a 20 s inter-stimulus interval while their behaviour was monitored at 500 fps. We also provided 250 ms trains of light pulses at 20 or 40 Hz. **B** Opsin expression in spinal motor neurons in a *Tg(mnx1:GAL4;UAS:CoChR-tdTomato)* larva at 5 dpf. Imaging field of view corresponds to black box in (**A**). A, anterior; D, dorsal; P, posterior; V, ventral. Scale bar 50 *μ*m. **C** Swim bouts elicited by a pulse train in *Tg(mnx1:GAL4;UAS:CoChR-tdTomato)* larvae (left). The control, opsin-negative larva (right), does not respond within 148 ms after stimulus onset. **D** Tail tracking, showing optogenetically-evoked swim bouts in a CoChR-expressing larva (bottom three rows) and a visually-evoked swim in a control opsin-negative larva (top). tbf, tail beat frequency. **E** Distribution of response latencies for all tail movements in opsin-expressing (red) and control opsin-negative larvae (grey). Dotted line indicates maximum latency (50 ms) for a response to be considered optogenetically-triggered. Control larvae exclusively show long latency responses. Each time bin corresponds to 25 ms. **F,L** Response probability of larvae expressing different opsins for single-pulse (**F**) or pulse-train (**L**) stimulation (mean ± SEM, across fish). **G–Q** Latency (**G,M**), bout duration (**H,N**), tail angle of the first half beat (θ_1_; **I,O**), number of cycles (**J,P**) and tail beat frequency (**K,Q**) for single-pulse (**G–K**) or pulse-train (**M–Q**) stimulation (mean ± SEM, across fish).

Optogenetically-evoked tail movements were triggered with short latency following light onset (8.3 ± 6.9 ms, mean ± SD) in opsin-expressing larvae only, whereas visually-evoked swim bouts occurred at much longer latency (316 ± 141 ms, mean ± SD) in both opsin-expressing larvae and control siblings (Figure 3E). We restricted our analyses to optogenetically-evoked movements, initiated within 50 ms of stimulus onset (corresponding to a minimum of the probability density distribution of latency; dotted line in Figure 3E). Optogenetically-evoked tail movements comprised a sequence of left-right alternating half beats, thereby resembling natural swim bouts (Figure 3C,D and Video 2,3). Response probability increased with irradiance (Figure 3F and Figure 3–figure supplement 1) and CoChR again elicited tail movements with the highest probability and shortest latency in response to blue light (96–100% at 0.63–2.55 mW/mm^2^; Figure 3F,G). Only the ChrimsonR line responded to red light (∼78% response probability at 1 mW/mm^2^; Figure 3F). Tail movements evoked by single light pulses typically had shorter duration and fewer cycles than visually-evoked swims (Figure 3H–K). However, longer movements (> 100 ms, 4–5 cycles) were often observed in response to single light pulses (see response to 2 ms pulse in Figure 3D and Video 2) indicating engagement of spinal central pattern generators. This may occur through recruitment of glutamatergic V2a interneurons connected to motor neurons via gap junctions (Song *et al*., 2016) and/or by proprioceptive feedback via cerebrospinal fluid-contacting neurons (Wyart *et al*., 2009; Fidelin *et al*., 2015; Bohm *et al*., 2016). Pulse train stimuli evoked swim bouts of longer duration, with swims in CoChR and ChrimsonR lines showing modest frequency-dependent modulation of cycle number (Figure 3L–Q).

### *In vivo* whole-cell recording of photocurrents in larval primary motor neurons

To calibrate photocurrents *in vivo*, we performed whole-cell voltage clamp recordings from single primary motor neurons (pMNs) in 5–6 dpf larvae (Figure 4A). Each opsin was stimulated with a wavelength close to its absorption peak (1–30 mW/mm^2^; Figure 4–figure supplement 1A). We recorded over 125 neurons, including control cells from opsin-negative animals, from which 86 cells were selected following strict criteria for recording quality (see Material and methods; N = 3–19 included cells per group; Figure 4–figure supplement 1B). Opsin-expressing pMNs displayed physiological properties, such as membrane resistance, resting membrane potential and cell capacitance, comparable to opsin-negative neurons (Figure 4B,C and Figure 4–figure supplement 1C,D). All cation channelrhodopsins induced inward currents upon light stimulation, which were not observed in opsin-negative pMNs (Figure 4D). Notably, CoChR and ChrimsonR generated the largest photocurrents (CoChR 475 ± 186 pA, mean ± SD, N = 8 cells, ChrimsonR 251 ± 73 pA, N = 7; Figure 4E). We did not observe significant irradiance-dependent modulation of photocurrent amplitude in any opsin line, likely due to the high range of irradiance we tested (Figure 4–figure supplement 1F). Photocurrent kinetics influence the temporal precision with which single action potentials can be evoked (Mattis *et al*., 2011). Therefore, we measured the photocurrent activation time (i.e. time to peak response from light onset), which results from the balance between activation and inactivation of the opsin, and deactivation time constant (i.e. the response decay time constant, τ_off_), which is determined by the rate of channel closure at light offset (Mattis *et al*., 2011; Schneider *et al*., 2015). Comparable activation times were observed across opsin lines (4–5 ms; Figure 4F). Deactivation time constants were more variable between opsins, with Chronos showing the fastest deactivation kinetics (4.3 ± 0.4 ms, N = 3 cells, mean ± SD) and the other opsins displaying similar time constants (12–20 ms; Figure 4G).

**Figure 4.**
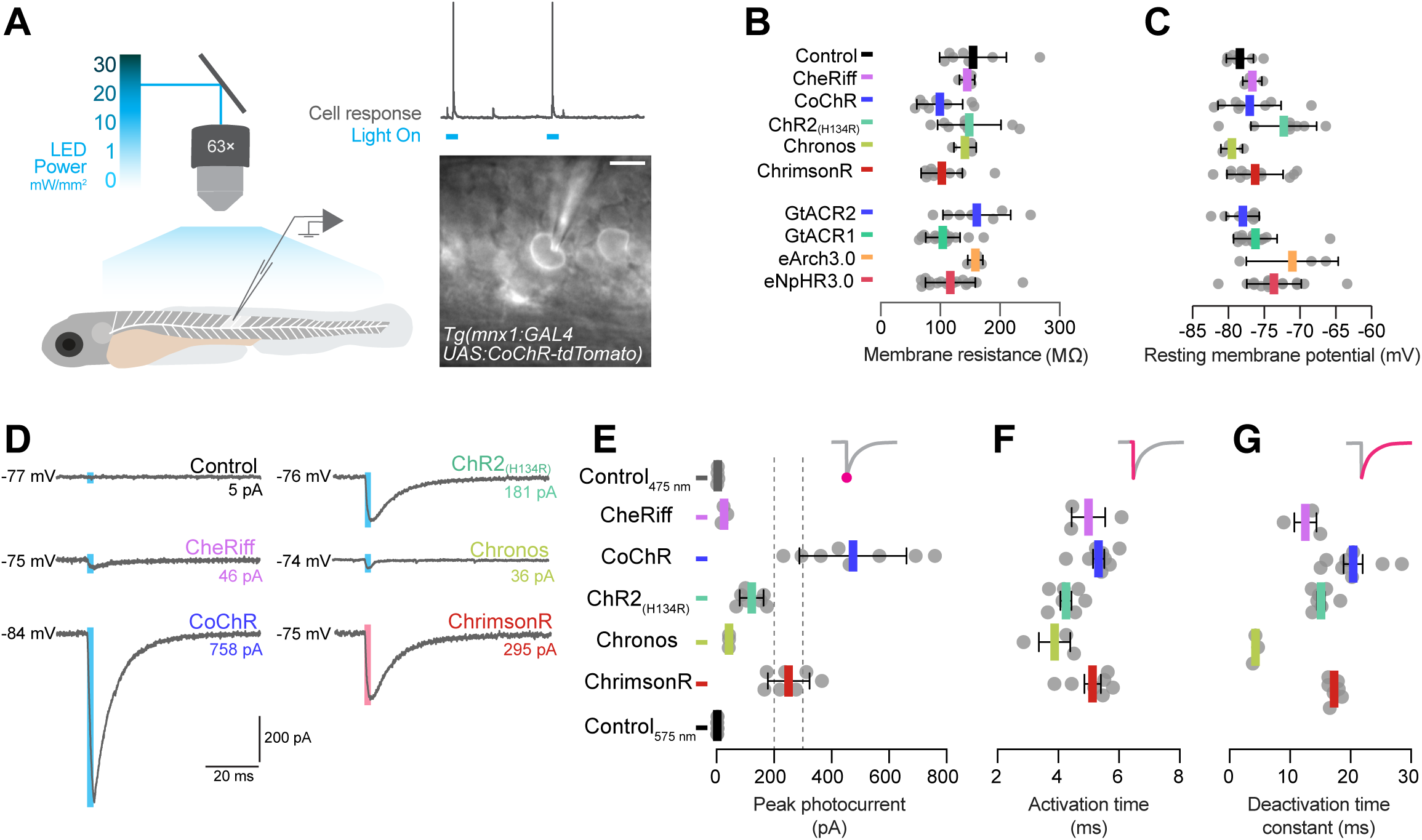
Electrophysiological recording of photocurrents in primary motor neurons. **A** Schematics of experimental setup for optogenetic stimulation with *in vivo* whole-cell patch clamp recordings. Image shows a patched primary motor neuron (pMN) expressing CoChR in a 6 dpf *Tg(mnx1:GAL4;UAS:CoChR-tdTomato)* larva. Scale bar 5 *μ*m. **B** Membrane resistance was not affected by opsin expression (mean ± SD, across cells). **C** Resting membrane potential was similar between opsin-expressing and control neurons (mean ± SD). **D** Examples of inward photocurrents in response to 5 ms light pulses (20 mW/mm^2^). **E** Peak photocurrent amplitude. CoChR and ChrimsonR induced the largest photocurrents (mean ± SEM, across cells). Dotted lines show range of pMN rheobase. Data is pooled across stimulus intensity (1–30 mW/mm^2^) but see Figure 4–figure supplement 1 for data at varying irradiance. **F** Photocurrent activation time was similar across opsins (mean ± SEM). **G** Chronos photocurrents had the fastest deactivation time constant, while CoChR and ChrimsonR showed similar deactivation kinetics (mean ± SEM).

### Optogenetic induction of spiking in larval pMNs

To investigate whether our cation channelrhodopsin lines can induce action potentials in pMNs, we performed *in vivo* current clamp recordings while providing single light pulses (1–5 ms duration). In all opsin lines, light stimulation induced voltage depolarisations, which were never observed in opsin-negative pMNs, and voltage responses above –30 mV were classified as spikes (Figure 5A).

**Figure 5.**
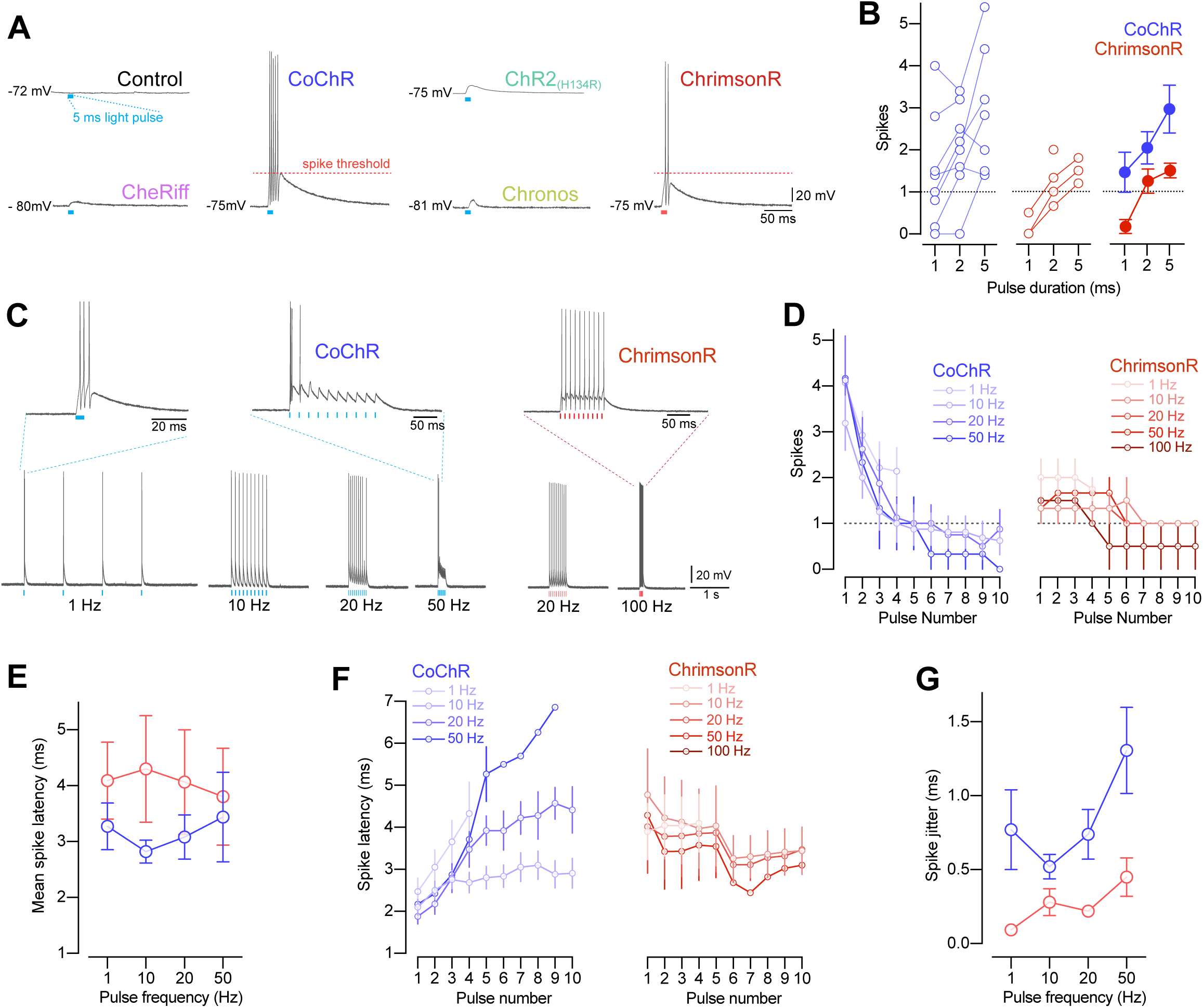
CoChR and ChrimsonR elicited spiking in primary motor neurons. **A** Example membrane depolarisations induced by 5 ms light pulses (20 mW/mm^2^). **B** Number of optogenetically-evoked spikes vs. pulse duration (across irradiance levels 1– 30 mW/mm^2^). Longer pulse duration induced more spikes in both CoChR- and ChrimsonR-expressing cells. Left plots show single neurons and right plot shows mean ± SEM across cells. **C** Example voltage responses from CoChR- and ChrimsonR-expressing cells upon pulse train stimulation (1–100 Hz, 2–5 ms pulse duration). **D** Number of spikes vs. pulse number within a train (mean ± SEM, across cells). In CoChR-expressing cells, the initial 3–4 pulses of the train induced bursts of 2–4 spikes. **E** Mean spike latency vs. pulse frequency (mean ± SEM). **F** Spike latency vs. pulse number (mean ± SEM). With increasing pulse frequency, CoChR-expressing cells showed progressively longer spike latency throughout the pulse train. **G** Spike jitter vs. pulse frequency (mean ± SEM). ChrimsonR-expressing cells showed lower spike jitter than CoChR-expressing cells.

CoChR and ChrimsonR were the only opsin lines capable of triggering spiking in this cell type (Figure 5A and Figure 5–figure supplement 1A–C), as expected from their peak photocurrents exceeding pMN rheobase (dotted lines in Figure 4E). Notably, 5 ms-long light pulses induced spikes in all CoChR-expressing neurons (N = 7 out of 7 cells at 3–30 mW/mm^2^), dropping to 88% of cells spiking with shorter pulses (Figure 5–figure supplement 1A). ChrimsonR was less effective than CoChR in inducing action potentials, with 36–38% of neurons spiking when using 2–5 ms-long pulses (2 ms, N = 4 out of 11; 5 ms, N = 3 out of 8 cells) and only 13% spiking with 1 ms-long pulses (N = 1 out of 8 cells). In both opsin lines, the number of evoked spikes increased with longer pulse duration (Figure 5B and Figure 5–figure supplement 1D).

For experiments aiming to replay physiological firing patterns, optogenetic actuators should be capable of inducing spike trains with millisecond precision and at biological firing frequencies. We thus tested the ability of CoChR and ChrimsonR to evoke pMN firing patterns across a range of frequencies (1–100 Hz; Figure 5C). pMNs can spike at high frequency (up to 300–500 Hz) (Menelaou and McLean, 2012), hence optogenetic induction of high-frequency firing should not be limited by cell intrinsic physiological properties, but rather by opsin properties and light stimulation parameters. To assess the fidelity of firing patterns at each stimulation frequency, we measured spike number per light pulse as well as spike latency and jitter (i.e. standard deviation of spike latency). ChrimsonR could induce firing up to the highest frequency tested (100 Hz), with each light pulse typically evoking a single spike (Figure 5C,D). CoChR generated spike bursts in response to the initial pulses of the train only and could not evoke spiking at stimulation frequencies higher than 50 Hz (Figure 5C,D). Overall, spikes were induced with short latency (3–4 ms mean latency) and low jitter (0.25–1.25 ms jitter) with both opsin lines (Figure 5E,G).

### Optogenetic suppression of coiling behaviour in embryos

Next, we tested the ability of our opsin lines to suppress spontaneous behaviour of zebrafish embryos (Saint-Amant and Drapeau, 1998; Warp *et al*., 2012; Mohamed *et al*., 2017; Bernal Sierra *et al*., 2018). We targeted expression of the anion-conducting channels GtACR1 and GtACR2 (Govorunova *et al*., 2015), the outward proton pump eArch3.0 (Mattis *et al*., 2011) and the inward chloride pump eNpHR3.0 (Gradinaru *et al*., 2010) to spinal cord neurons using the *Tg(s1020t:GAL4)* transgene (Scott *et al*., 2007) and examined changes in spontaneous coiling behaviour in response to light (Figure 6A– D and Video 4). In opsin-expressing embryos (24–27 hpf), light exposure led to a suppression of coiling behaviour that was followed by a synchronised restart at light offset (Figure 6D,E and Figure 6–figure supplement 1; N = 91 ± 16 fish per group, mean ± SD), as previously reported (Warp *et al*., 2012; Mohamed *et al*., 2017). As expected from behaviour with *Tg(isl2b:GAL4)* embryos (Figure 2F,G), GtACR activation in spinal neurons occasionally induced movements in the initial 1–2 s following light onset (black arrows in Figure 6D,E), a phenomenon that was not observed with Cl^−^/H^+^ pumps.

**Figure 6.**
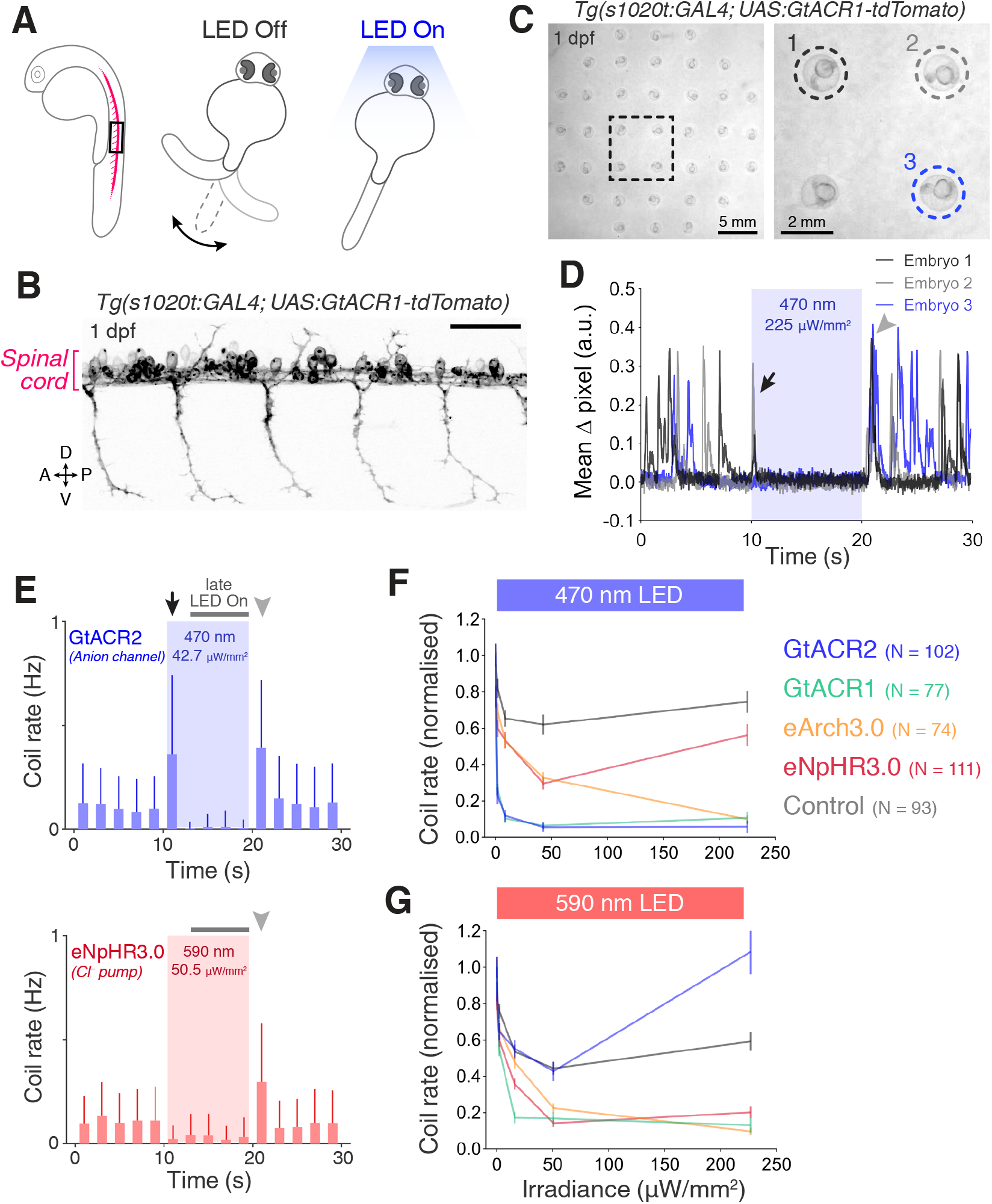
Optogenetic suppression of coiling behaviour in embryos. **A** Schematic of the behavioural assay. **B** Opsin expression in spinal motor neurons and interneurons in a *Tg(s1020t:GAL4;UAS:GtACR1-tdTomato)* embryo at 1 dpf. Imaging field of view corresponds to black box in (**A**). A, anterior; D, dorsal; P, posterior; V, ventral. Scale bar 50 *μ*m. **C** Camera field of view showing *Tg(s1020t:GAL4;UAS:GtACR1-tdTomato)* embryos positioned in individual agarose wells. Behaviour was monitored at 50 frames per second across multiple embryos (24–27 hpf; N = 91 ± 16 fish per group, mean ± SD) subjected to 10 s light periods (470 or 590 nm, 0–227 *μ*W/mm^2^) with a 50 s inter-stimulus interval. **D** Tracking of coiling behaviour (mean ΔPixel from 3 trials) for the 3 embryos shown in (**C**). Black arrow indicates movements at light onset, whereas grey arrowhead indicates synchronised restart of coiling behaviour following light offset. **E** Optogenetically-induced changes in coil rate (mean + SD, across fish) in embryos expressing the anion channelrhodopsin GtACR1 (N = 77 embryos, top) or the Cl^−^ pump eNpHR3.0 (N = 111 embryos, bottom). Horizontal dark grey bars indicate the ’late LED On’ period. Each time bin corresponds to 2 s. **F,G** Normalised coil rate during the ’late LED On’ period in embryos expressing different opsins (mean ± SEM, across fish). Control opsin-negative siblings were subjected to the same light stimuli.

Given these two effects, changes in coil rate were separately quantified for the initial 2 s (Figure 6– figure supplement 2) and subsequent 8 s period of light exposure (‘late LED ON’; grey horizontal bars in Figure 6E).

All opsin lines suppressed coiling behaviour during the ‘late LED ON’ period (Figure 6F,G). This was likely a result of distinct mechanisms: hyperpolarisation with Cl^−^/H^+^ pumps versus depolarisation block with anion channelrhodopsins (see below and Discussion). As previously observed (Friedmann *et al*., 2015), light also decreased coiling in control opsin-negative embryos, yet to a significantly lesser degree than in opsin-expressing animals (Figure 6F,G). GtACRs achieved the strongest suppression of coil rate using blue light (90–95% decrease at 8.4–225 *μ*W/mm^2^; Figure 6F). With amber light, GtACR1, eArch3.0 and eNpHR3.0 showed comparable suppression (80–90% decrease at 50.5– 227 *μ*W/mm^2^), with GtACR1 achieving ∼83% decrease in coil rate even at low irradiance (15.9 *μ*W/mm^2^; Figure 6G).

### Optogenetic suppression of swimming in larvae

To compare the efficacy of our opsin lines to suppress behaviour in larvae, we targeted opsin expression to spinal motor neurons and interneurons using *Tg(s1020t:GAL4)*, as above, and examined changes in spontaneous swimming behaviour of 6 dpf animals in response to 10 s-long light pulses (Figure 7A–C and Video 5; N = 25 ± 9 fish per group, mean ± SD).

**Figure 7.**
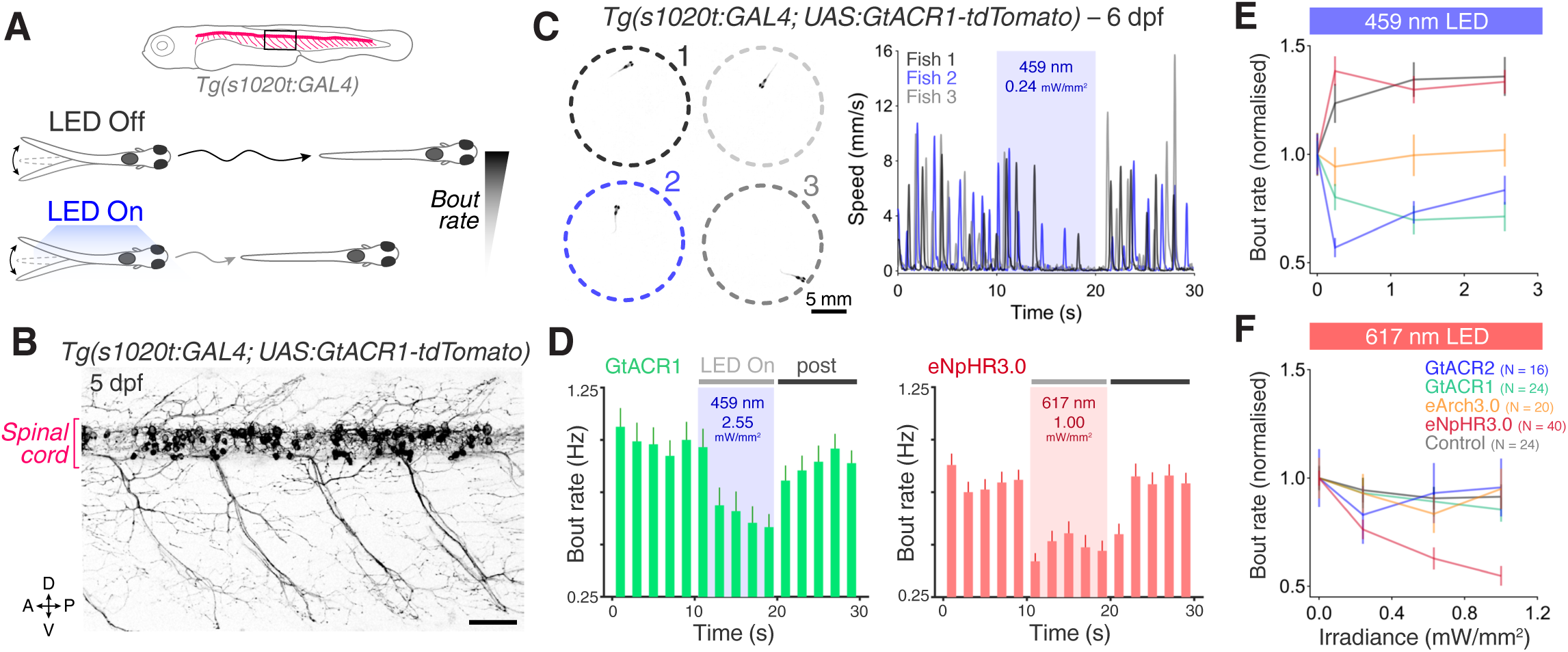
Optogenetic suppression of swimming in larvae. **A** Schematic of behavioural assay. **B** Opsin expression in spinal motor neurons and interneurons in a *Tg(s1020t:GAL4;UAS:GtACR1-tdTomato)* larva at 5 dpf. Imaging field of view corresponds to black box in (**A**). A, anterior; D, dorsal; P, posterior; V, ventral. Scale bar 50 *μ*m. **C** *Tg(s1020t:GAL4;UAS:GtACR1-tdTomato)* larvae were positioned in individual agarose wells (left) and instantaneous swim speed was monitored by centroid tracking (right) at 50 fps (6 dpf; N = 25 ± 9 fish per group, mean ± SD). 10 s light periods were delivered (459 or 617 nm, 0–2.55 mW/mm^2^) with a 50 s inter-stimulus interval. **D** Optogenetically-induced changes in bout rate (mean + SEM, across fish) in *Tg(s1020t:GAL4)* larvae expressing GtACR1 (N = 24 larvae, left) or eNpHR3.0 (N = 40 larvae, right). Horizontal grey bars indicate the time windows used to quantify behavioural changes. Each time bin corresponds to 2 s. **E,F** Normalised bout rate during the ‘LED On’ period in larvae expressing different opsins (mean ± SEM, across fish) and in control, opsin-negative, siblings.

GtACR1, GtACR2 and eArch3.0 reduced swim bout rate relative to control larvae in response to blue light, with GtACRs achieving the greatest suppression (20–45% decrease; Figure 7D,E). Consistent with a previous report (Andalman *et al*., 2019), opsin-negative larvae showed a 20–30% increase in bout rate during illumination with blue light (Figure 7E and Figure 7–supplement 1), while no increase was observed with red light (Figure 7F). Using red light, only eNpHR3.0 could reduce bout rate and suppression increased with higher irradiance (45% decrease at 1 mW/mm^2^; Figure 7F). No increase in bout rate was found in larvae expressing anion channelrhodopsins even when analysis was restricted to the initial 2 s of the light period (Figure 7–figure supplement 2A), suggesting GtACRs do not induce excitatory effects at larval stages. Opsin activation did not affect bout speed (Figure 7–figure supplement 2B). By contrast, using the *Tg(mnx1:GAL4)* transgene to drive opsin expression in motor neurons resulted in a decrease in bout speed (∼20% reduction), but not bout rate (Figure 7–figure supplement 3,4).

### Photocurrents induced by anion channelrhodopsins and chloride/proton pumps

To analyse the physiological effects induced by anion channelrhodopsins and Cl^−^/H^+^ pumps, we measured their photocurrents through *in vivo* voltage clamp recordings from larval pMNs (5–6 dpf). Since anion channelrhodopsin function depends on chloride homeostasis (Figure 8A) (Govorunova *et al*., 2015) and chloride reversal potential (ECl) is known to change over development (Ben-Ari, 2002; Reynolds *et al*., 2008; Zhang *et al*., 2010), we recorded GtACR1 photocurrents using two intracellular solutions: one mimicking ECl in embryonic neurons (–50 mV) (Saint-Amant and Drapeau, 2003) and the second approximating intracellular chloride concentration in more mature, larval neurons (ECl = –70 mV, see Materials and methods). Inspection of I-V curves for GtACR1 photocurrents showed that, in both solutions, currents reversed with a positive 5–10 mV shift relative to ECl (Figure 8–supplement 1A,B), as previously observed (Govorunova *et al*., 2015) and within the expected error margin given our access resistance (Figure 4–figure supplement 1C; estimated voltage error for ECl–50 mV solution, 4.6 ± 6.4 mV, mean ± SD, N = 5 cells; ECl–70 mV solution, 1.2 ± 1.3 mV, N = 3). This suggests that GtACR1 photocurrents were primarily driven by chloride ions, as expected (Govorunova *et al*., 2015). The other opsin lines were tested using the ECl–50 mV solution only. Neurons were stimulated with light (1 s-long pulse) at a holding potential matching their measured resting membrane potential (Figure 4C).

**Figure 8.**
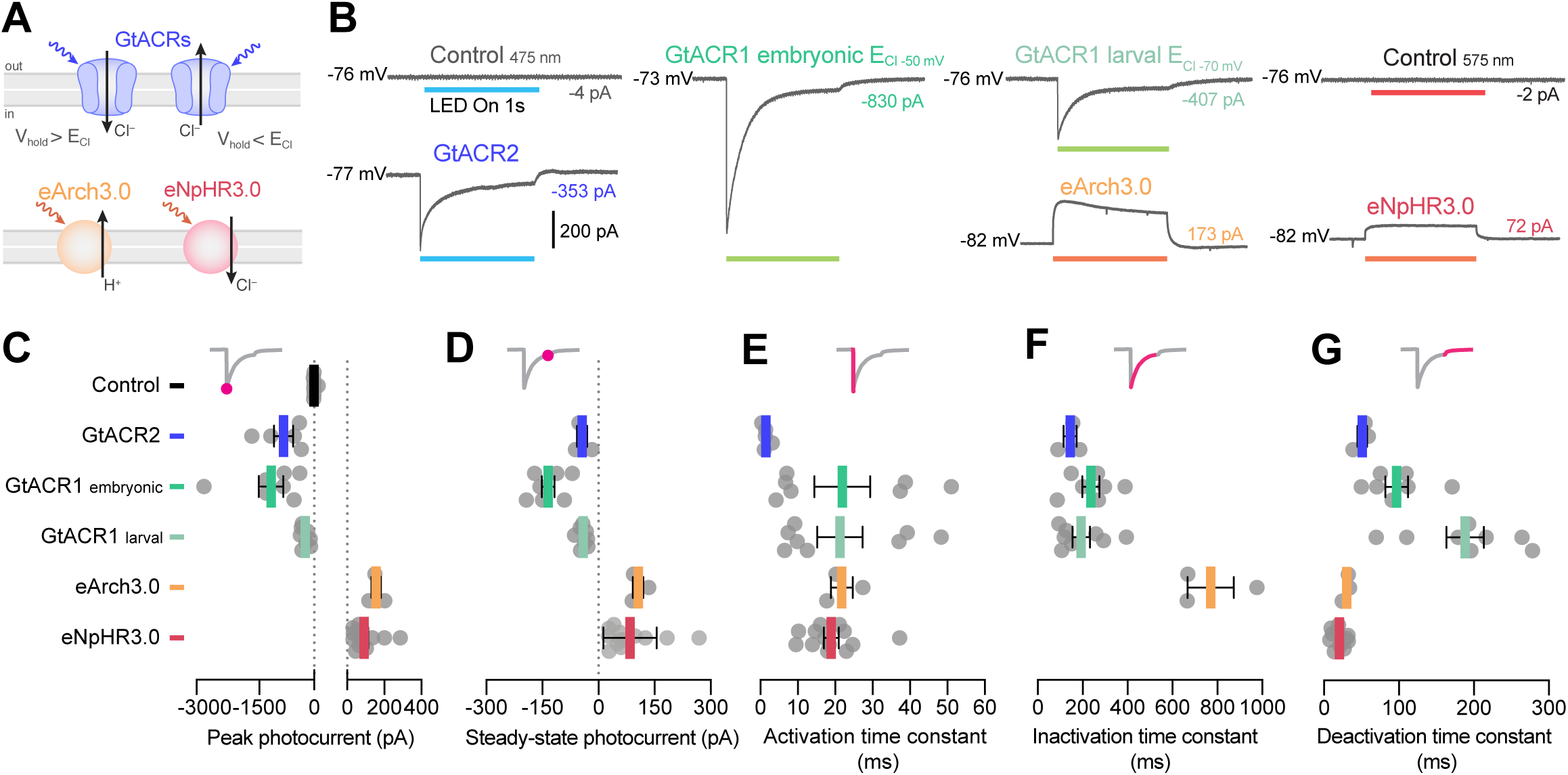
Photocurrents induced by anion channelrhodopsins and chloride/proton pumps. **A** Action of anion channelrhodopsins (top) and Cl^−^/H^+^ pumps (bottom). For anion channelrhodopsins, photocurrent magnitude and direction depend on chloride reversal potential (ECl) and holding potential (V_hold_), while Cl^−^/H^+^ pumps always induce outward currents. **B** Example photocurrents in response to a 1 s light exposure (20 mW/mm^2^). **C,D** Photocurrent peak (**C**) and steady-state (**D**) amplitude (mean ± SEM, across cells). GtACRs induced larger photocurrents than Cl^−^/H^+^ pumps. **E–G** Photocurrent activation (**E**), inactivation (**F**) and deactivation (**G**) time constants (mean ± SEM). Photocurrents induced by Cl^−^/H^+^ pumps showed minimal inactivation and faster deactivation kinetics than GtACRs. eNpHR3.0 photocurrents did not inactivate hence no inactivation time constant was computed.

Anion channelrhodopsins induced inward, ‘depolarising’ photocurrents (as expected from the combination of ECl and holding potential), while Cl^−^/H^+^ pumps generated outward, ‘hyperpolarising’ currents (Figure 8B). All opsins except eNpHR3.0 showed bi-phasic photocurrent responses composed of a fast activation followed by a slow inactivation (Figure 8B), likely due to a fraction of the opsin population transitioning to an inactive state (Chow *et al*., 2010; Mattis *et al*., 2011; Schneider *et al*., 2015). We measured both the peak photocurrent (Figure 8C) as well as the steady-state current during the last 5 ms of the light period (Figure 8D). GtACRs induced photocurrents with peak amplitude 3–10 times larger than those generated by Cl^−^/H^+^ pumps (Figure 8C), while steady-state currents were similar across opsins (Figure 8D). Some degree of irradiance-dependent modulation of photocurrents was observed, primarily in peak amplitude (Figure 8–supplement 2C–To characterise photocurrent kinetics, we computed activation, inactivation (or !des) and deactivation time constants (Mattis *et al*., 2011). GtACR photocurrents had the fastest activation kinetics (∼1 ms at 30 mW/mm^2^; Figure 8E and Figure 8–figure supplement 2F). However, deactivation kinetics of Cl^−^/H^+^ pumps were 2–10 times faster than those induced by GtACRs (14– 22 ms eNpHR3.0, 27–37 ms eArch3.0; Figure 8G and Figure 8–figure supplement 2H) and showed little inactivation (600–1000 ms eArch3.0; Figure 8F and Figure 8–figure supplement 2G).

### Optogenetic inhibition of pMN spiking

To investigate the ability of anion channelrhodopsins and Cl^−^/H^+^ pumps to suppress neural activity, we recorded pMNs in current clamp mode. In control opsin-negative neurons, light delivery (1 s) induced negligible voltage deflections (Figure 9A). By contrast, anion channelrhodopsins generated membrane depolarisation towards ECl while the Cl^−^/H^+^ pumps hyperpolarised the cell (Figure 9A), in accordance with recorded photocurrents. The absolute peak amplitude of voltage deflections was comparable between opsin lines (10–25 mV), with 10–40% decrease between peak and steady-state responses in all cases except eNpHR3.0, which generated stable hyperpolarisation (Figure 9B,C and Figure 9–figure supplement 1A,B). In a subset of GtACR1-(N = 4 out of 7) and GtACR2-expressing neurons (N = 2 out of 6), spiking was induced at light onset when using the ECl_–50 mV_ solution (Figure 9A; GtACR1 6.7 ± 7.1 spikes; GtACR2 1.5 ± 0.7, mean ± SD). This is consistent with the movements evoked at light onset in young, 1 dpf embryos expressing GtACRs (Figure 2 and 6). The kinetics of voltage decay to baseline following light offset matched those of recorded photocurrents (Figure 9D and Figure 9–figure supplement 1C).

**Figure 9.**
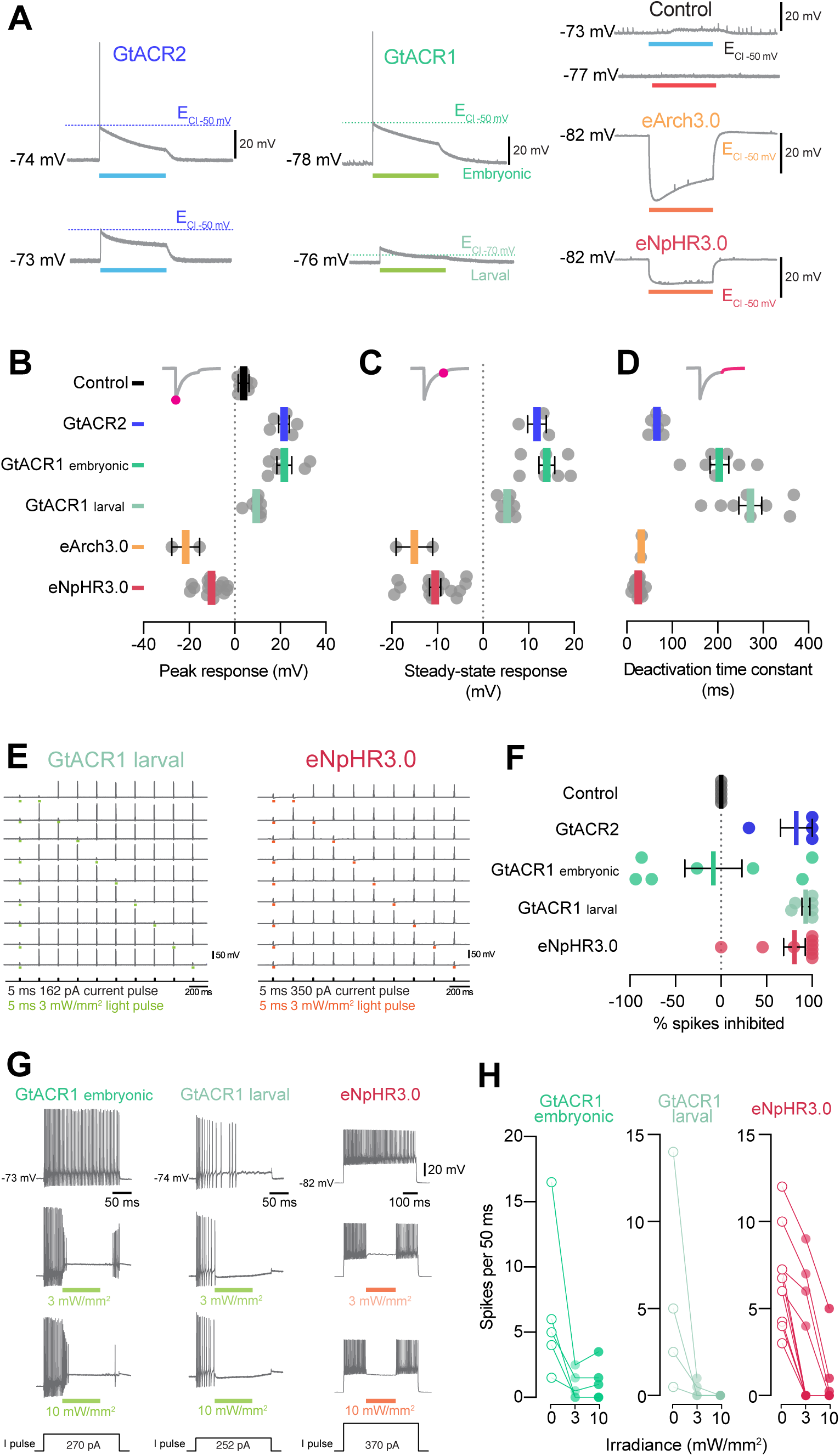
GtACRs and eNpHR3.0 effectively inhibited spiking. **A** Example voltage deflections induced by anion channelrhodopsins and Cl^−^/H^+^ pumps in response to a 1 s light pulse (20 mW/mm^2^). **B–D** Peak (**B**) and steady-state (**C**) responses and deactivation time constant (**D**) of voltage deflections. All opsins induced similar absolute voltage changes. Anion channelrhodopsins generated depolarisation with both intracellular solutions while Cl^−^/H^+^ pumps generated hyperpolarisation. **E** Example recordings demonstrating inhibition of single spikes in GtACR1- and eNpHR3.0-expressing cells with 5 ms light pulses (3 mW/mm^2^). **F** Fraction of spikes that were optogenetically inhibited (mean ± SEM, across cells). All opsins achieved high suppression efficacy, but GtACR1 induced additional spikes upon light delivery with the embryonic intracellular solution. **G** Example recordings demonstrating inhibition of sustained spiking in GtACR1- and eNpHR3.0-expressing cells. **H** Quantification of suppression using protocol illustrated in G. Number of spikes per 50 ms during light delivery (0–10 mW/mm^2^) is plotted against irradiance. GtACR1 and eNpHR3.0 inhibited tonic spiking with similar efficacy (mean ± SEM).

Next, we compared the utility of our opsin lines to inhibit pMN firing. First, we induced larval pMNs to fire at 5 Hz by injecting pulses of depolarising current (5 ms, 1.2–1.5× rheobase) and simultaneously delivered 5 ms light pulses to inhibit selected spikes (Figure 9E). We found that GtACRs and eNpHR3.0 could effectively inhibit spikes (80–95% suppression), while light pulses did not alter firing in opsin-negative neurons (Figure 9F). In agreement with our current clamp recordings, a subset of GtACR1-expressing neurons (N = 4 out of 7) tested in the embryonic ECl–50 mV solution failed to suppress spikes and instead induced extra action potentials in response to light pulses, resulting in a negative spike inhibition efficacy (Figure 9F). Data from eArch3.0-expressing neurons could not be collected due to degradation in the quality of recordings or cells becoming highly depolarised (i.e. resting membrane potential > –50 mV) by the later stages of the protocol, suggesting that repeated eArch3.0 activation may alter electrical properties of neurons (Williams *et al*., 2019).

Lastly, we asked whether we could inhibit firing over periods of tens to hundreds of milliseconds. We injected long pulses of depolarising current (200–800 ms) to elicit tonic pMN firing, and simultaneously provided shorter light pulses (50–200 ms; 3–10 mW/mm^2^) in the middle of the spike train (Figure 9G). Both GtACR1 and eNpHR3.0 successfully inhibited spiking during the light pulse, with complete suppression in 60–100% of cells at 10 mW/mm^2^ irradiance (Figure 9G,H). Notably, GtACR1 could inhibit tonic spiking even when using the embryonic ECl_–50 mV_ solution (Figure 9G,H), consistent with the suppression of coiling behaviour upon prolonged illumination of GtACR-expressing embryos (Figure 6).

## Discussion

In this study, we generated a set of stable transgenic lines for GAL4/UAS-mediated opsin expression in zebrafish and evaluated their efficacy in controlling neural activity *in vivo.* High-throughput behavioural assays and whole-cell electrophysiological recordings provided complementary insights to guide tool selection. Behavioural assays enabled efficient evaluation of opsin lines in various sensory and motor cell types and revealed developmental stage-specific effects in intact neural populations. Electrophysiological recordings from single motor neurons afforded quantification of photocurrents and systematic evaluation of the ability of these optogenetic tools to elicit or silence activity at single action potential resolution.

### An *in vivo* platform for opsin tool selection

The selection of optogenetic actuators should be based on their ability to reliably control neural activity *in vivo.* While previous efforts compared opsin efficacy using transient expression strategies [e.g. viral or plasmid injections, see Mattis *et al*. (2011) and Introduction], here we calibrated opsin effects in stable transgenic lines, which offer more reproducible expression across experiments and laboratories (Kikuta and Kawakami, 2009; Yizhar *et al*., 2011). Overall, there was good qualitative agreement between behavioural and electrophysiological results, with efficacy in behavioural assays (even with transient expression) largely predicting rank order in photocurrent amplitudes. This illustrates the utility of high-throughput behavioural assays for rapid evaluation and selection of expression constructs prior to more time-consuming generation and characterisation of stable lines and electrophysiological calibration. We observed broad variation in efficacy across lines, likely attributable to differences in both the intrinsic properties of the opsin as well as variation in expression and membrane targeting. Membrane trafficking can also be influenced by the fluorescent protein fused to the actuator (Arrenberg *et al*., 2009). In our hands, we observed better expression with the tdTomato fusion reported here than with previous attempts using a tagRFP fusion protein. In the future, expression might be further improved through codon optimisation (Horstick *et al*., 2015), trafficking-enhancing sequences (Gradinaru *et al*., 2010; Mattis *et al*., 2011), alternative expression targeting systems (Luo *et al*., 2008; Sjulson *et al*., 2016) and optimisation of the fluorescent reporter protein.

Behavioural and electrophysiological readouts complemented one another and enriched the interpretation of our results. Electrophysiological recordings in a defined cell type allowed direct and comparative calibration of photocurrents. Although several opsin lines did not evoke action potentials in low-input-resistance pMNs, behavioural assays showed that all lines induced tail movements in larvae. This is likely due to recruitment of secondary motor neurons labelled by the *Tg(mnx1:GAL4)* transgene, which have higher input resistance (Menelaou and McLean, 2012). Behavioural assays at multiple ages revealed that anion channelrhodopsins can excite neurons in 1 dpf embryos which was corroborated by making whole-cell recordings using a patch solution reproducing the high intracellular chloride concentration observed in embryonic neurons (Reynolds *et al*., 2008; Zhang *et al*., 2010).

Overall, our platform enables efficient selection and calibration of optogenetic tools for *in vivo* neuroscience. It also enables opsin-specific optimisation of light delivery (i.e. wavelength, pulse duration, frequency and intensity). For example, we found that equivalent stimulation regimes produced different rates of spiking adaptation that impacted the ability to control high-frequency firing, depending on the specific line in question.

### Robust and precise optogenetic induction of spiking

Which opsin lines are best suited for reliable neural activation? Photocurrent amplitude, measured in pMNs, was predictive of the ability of opsin lines to induce behaviour via activation of distinct cell types at both larval and embryonic stages (CoChR > ChrimsonR > ChR2_(H134R)_ > Chronos ≥ CheRiff). The CoChR and ChrimsonR lines showed the highest expression levels among cation channelrhodopsins and were the only lines capable of inducing action potentials in pMNs, consistent with their photocurrent amplitudes exceeding pMN rheobase. Notably, CoChR evoked spikes in all pMNs tested and triggered behaviour with maximal response probability in larvae at irradiance levels as low as 0.63 mW/mm^2^.

Where precise control of a cell’s firing pattern is desired, electrophysiological calibration is essential to tune stimulation parameters for a specific opsin/cell-type combination. Our data indicate that long light pulses (2–5 ms) can lead to spike bursts and substantial firing rate adaptation during high-frequency stimulation, likely a result of plateau potentials and inactivation of voltage-gated sodium channels. Thus, although the CoChR line produced large-amplitude photocurrents and was highly efficient in evoking spikes, it was also prone to burst firing, which compromised spiking entrainment with high-frequency stimulations. Therefore, short light pulses (< 2 ms) are better suited for inducing high-frequency firing patterns with millisecond precision when using CoChR.

### Excitatory effects of anion channelrhodopsins

Anion channelrhodopsins induced movements at light onset in 1 dpf embryos as well as transient spiking in pMNs when using an intracellular solution that mimicked the high ECl (–50 mV) of immature neurons. This is consistent with GtACRs functioning as a light-gated chloride conductance (Govorunova *et al*., 2015). The transient nature of spiking and motor activity might be due to the initial large inward photocurrent depolarising neurons above spiking threshold, while the subsequent smaller inactivating current would lead to depolarisation block by clamping membrane potential close to ECl. Transient induction of action potentials with GtACRs has also been observed in rat cortical pyramidal neurons in brain slices (Malyshev *et al*., 2017) as well as cultured hippocampal neurons (Mahn *et al*., 2018) and has been attributed to antidromic spiking resulting from a positively shifted ECl in the axon (Mahn *et al*., 2016; Mahn *et al*., 2018). In light of this, the use of GtACRs in immature neurons or subcellular structures should be carefully calibrated and use of Cl^−^/H^+^ pumps may be preferable.

### Precise optogenetic inhibition of neural activity

To accurately suppress action potentials, opsin tools must be carefully selected with consideration for developmental stage and ECl-dependent effects as well as photocurrent kinetics. GtACRs generated large photocurrents with fast activation kinetics, which can explain why GtACR1 was effective in inhibiting single action potentials with short light pulses in larval pMNs. Cl^−^/H^+^ pump photocurrents instead showed fast deactivation kinetics, which allowed eNpHR3.0-expressing neurons to rapidly resume spiking at light offset. Differences in photocurrent kinetics between opsin classes – i.e. channels vs. pumps – may thus differentially affect the temporal resolution of activity inhibition and recovery, respectively. The combined behavioural and electrophysiological approach can be extended in the future to optogenetic silencers based on K^+^ channel activation, such as the recently introduced PAC-K (Bernal Sierra *et al*., 2018).

In conclusion, our calibrated optogenetic toolkit and associated methodology provide an *in vivo* platform for designing controlled optogenetic experiments and benchmarking novel opsins.

## Acknowledgements

The authors thank members of the Bianco lab and Wyart lab for helpful discussions. We thank staff from the UCL and ICM PHENOZ fish facilities for fish care and husbandry (UCL: Carole Wilson and team; ICM: Sophie Nunes-Figueiredo, Bogdan Buzurin, and Monica Dicu), Irene Arnold-Ammer and Enrico Kühn (MPI of Neurobiology) assisted with generation of transgenic lines. We also acknowledge support from the ICM electrophysiology core facility (CELIS-ePhys). P.A. was supported by a Sir Henry Wellcome Postdoctoral Fellowship (204708/Z/16/Z). A.D. was supported by a Marie Curie Incoming International Fellowship (H2020-MSCA-IF-2016 Project #752199). F.K. was supported by a HFSP long-term fellowship (LT393/2010). Generation of opsins in H. B.’s laboratory was supported by the Max Planck Society (H. B. and F. K.) and the DFG (SPP1926 Next-Generation Optogenetics). A Sir Henry Dale Fellowship from the Royal Society & Wellcome Trust (101195/Z/13/Z) and a UCL Excellence Fellowship were awarded to I.H.B. The work in C.W.’s lab was funded by Human Frontier Science Program (HFSP) Research Grant (RGP063-2018) and the New York Stem Cell Foundation (NYSCF-R-NI39). The work on electrophysiological calibration of opsins performed in ICM has also received funding from the ICM foundation, the program ‘Investissements d’avenir’ ANR-10-IAIHU-06 (Big Brain Theory ICM Program), ANR-11-INBS-0011 (NeurATRIS: Translational Research Infrastructure for Biotherapies in Neurosciences).

## Author Contributions

Conceptualisation: PA, AD, CD, IB, CW Methodology: PA, AD, CD Software: PA, AD, IB, CW Validation: PA, AD Formal analysis: PA, AD Investigation: PA, AD, CD, HM, KL, TH Resources: PA, AD, CD, FK, HB, IB, CW Data curation: PA, AD Writing—original draft: PA, AD, CW, IB Writing—review and editing: PA, AD, FK, HB, IB, CW Visualisation: PA, AD Funding acquisition: PA, AD, FK, HB, IB, CW Supervision: HB, IB, CW Project administration: CW, IB

## Materials and methods

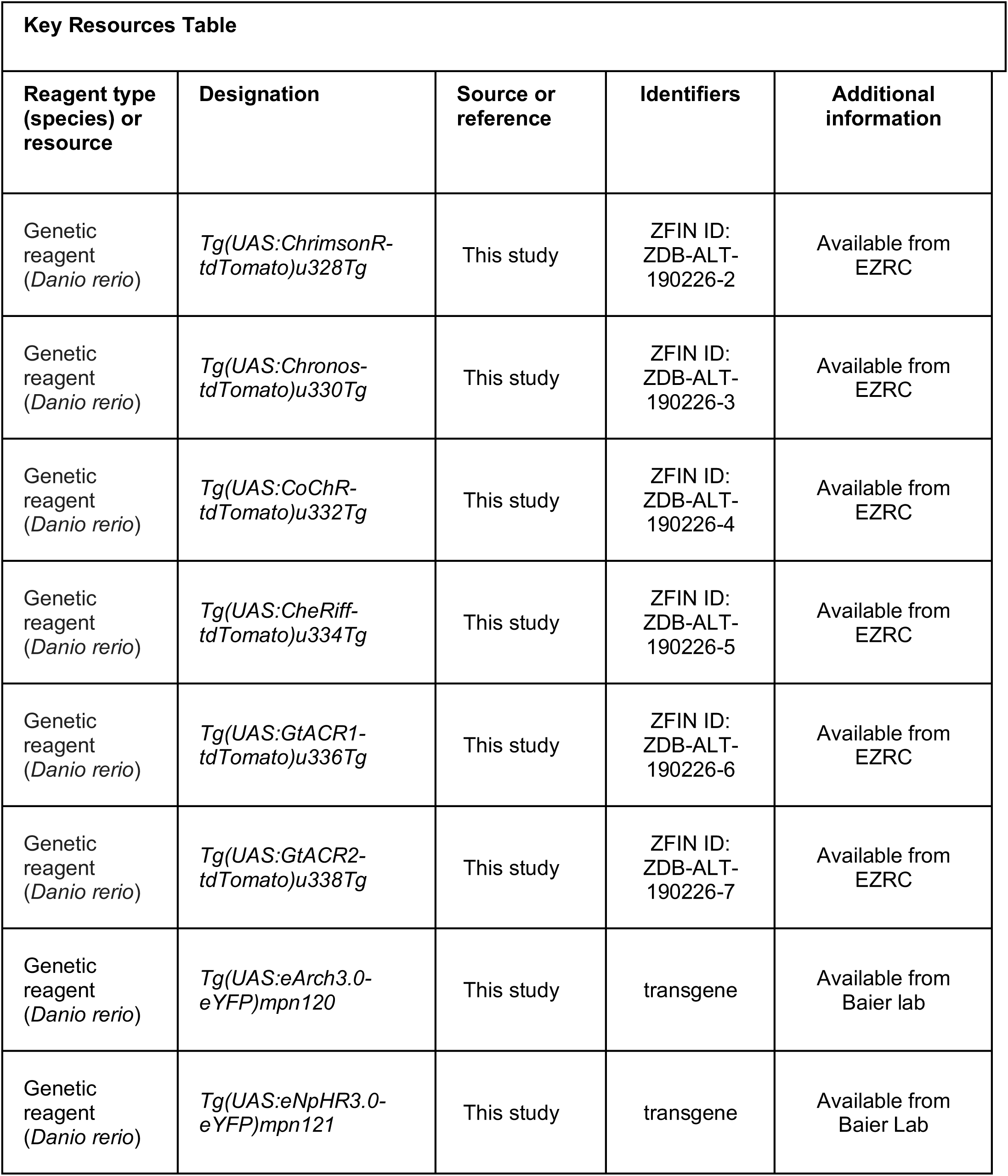
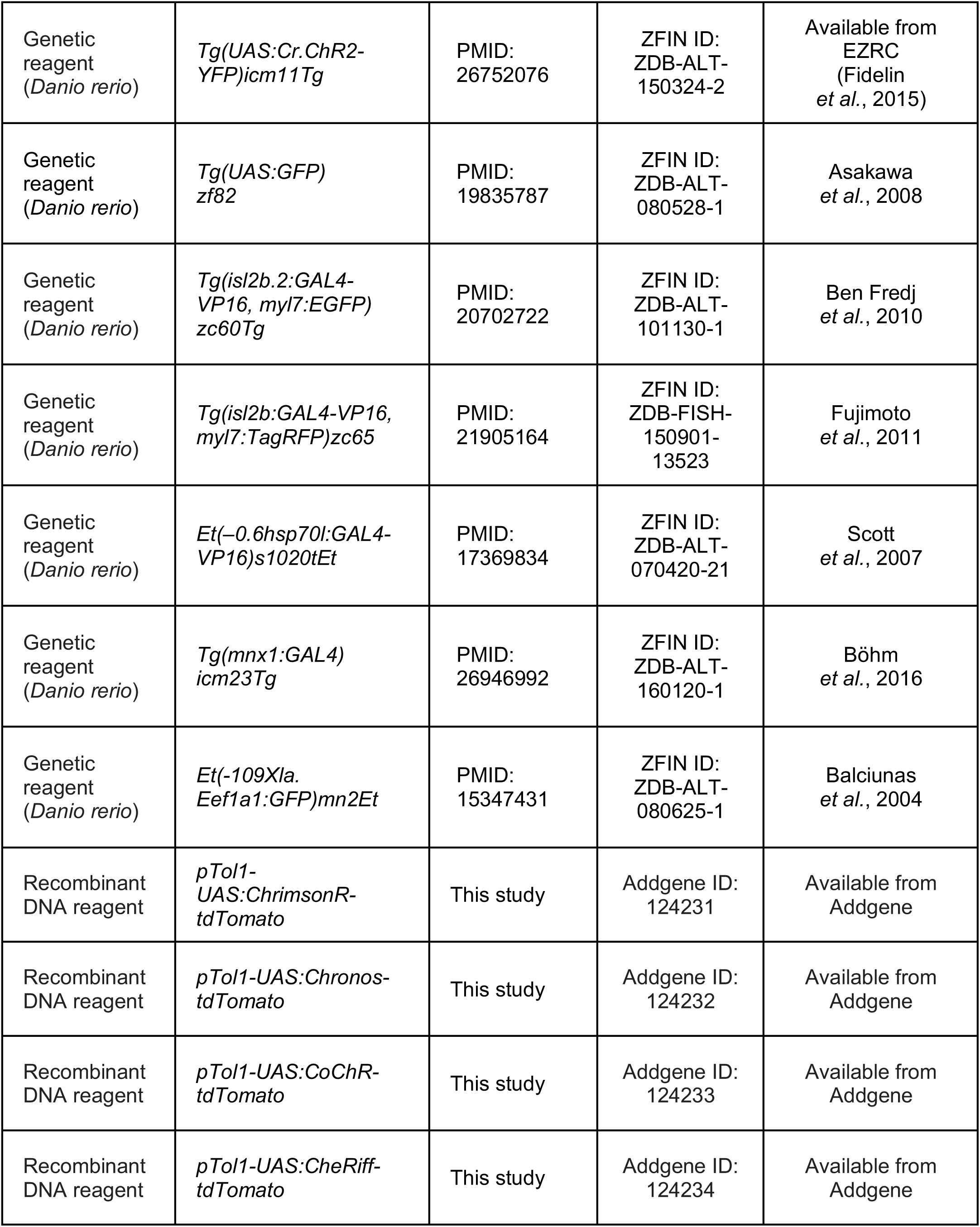
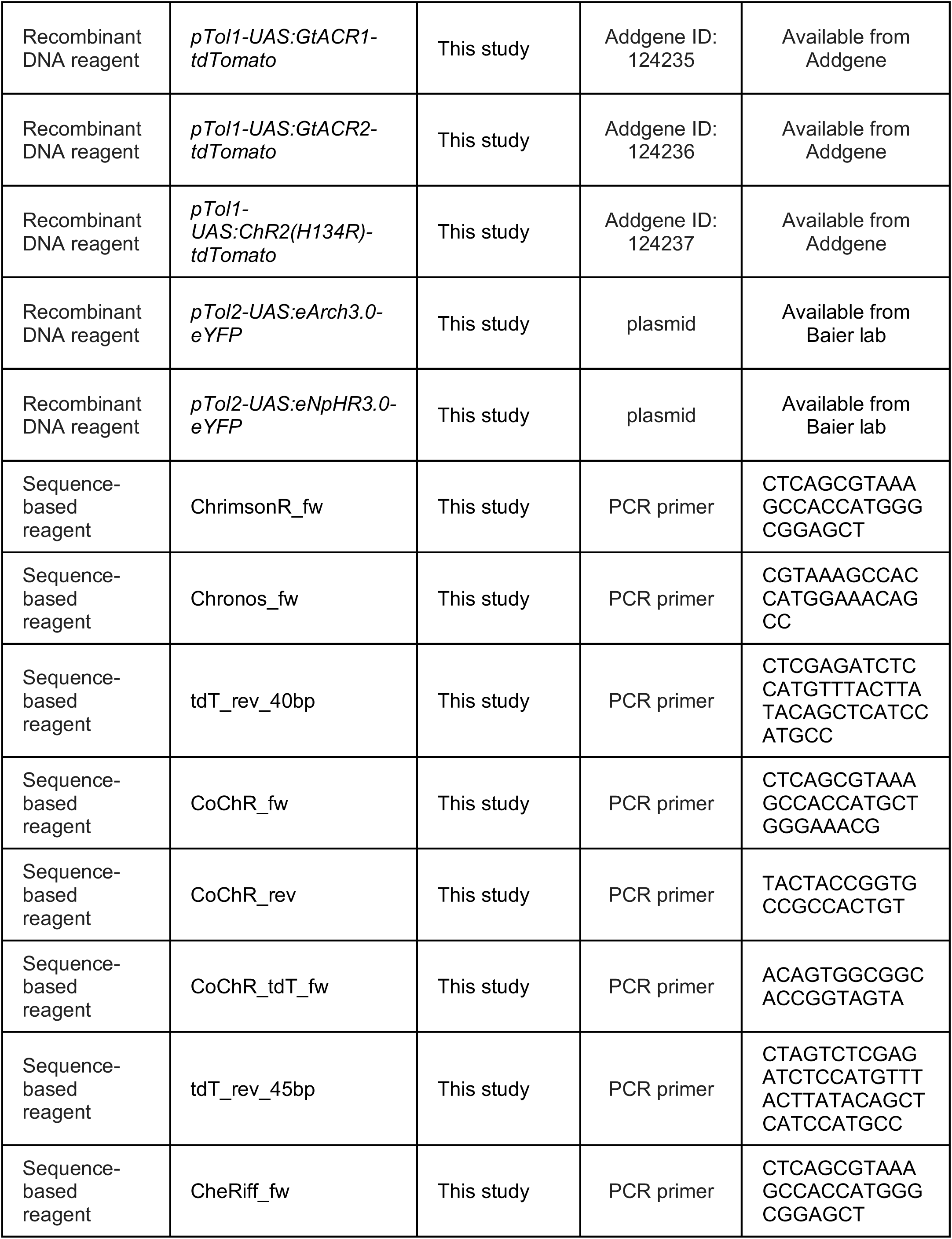
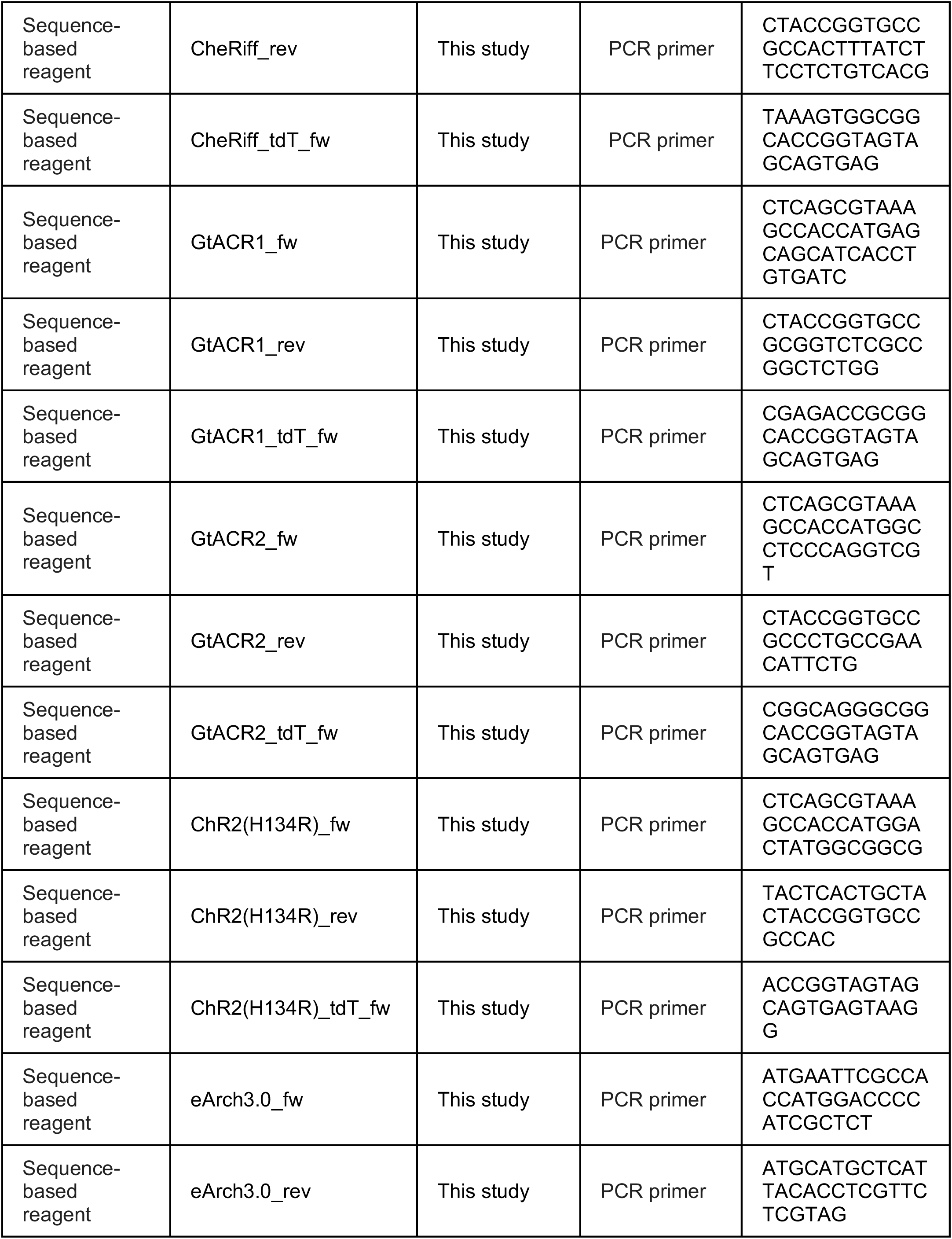
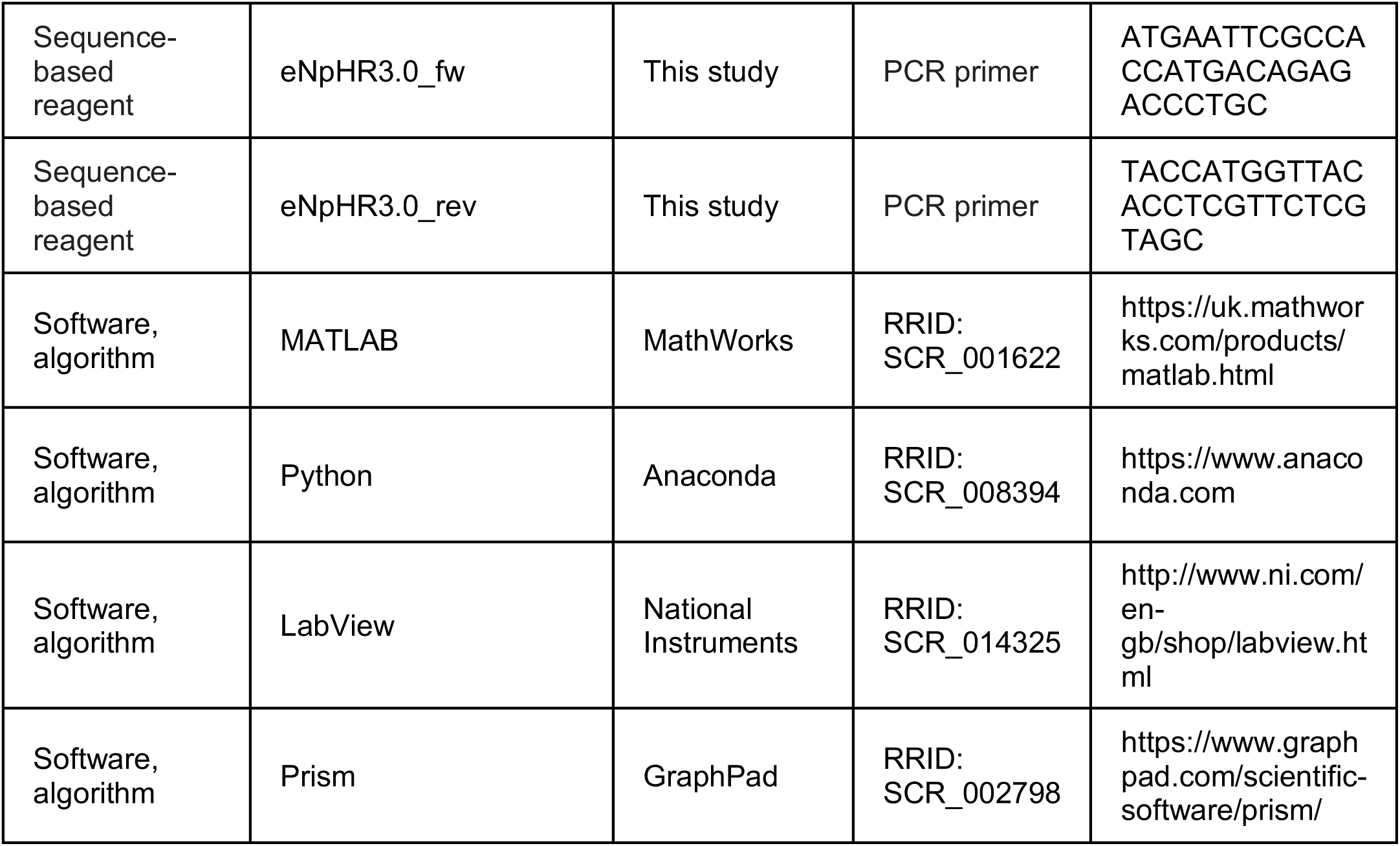

### Experimental model

Animals were reared on a 14/10 h light/dark cycle at 28.5°C. For all experiments, we used zebrafish (*Danio rerio*) embryos and larvae homozygous for the *mitfa^w2^* skin-pigmentation mutation (Lister *et al*., 1999). All larvae used for behavioural assays were fed *Paramecia* from 4 dpf onward. Animal handling and experimental procedures were approved by the UCL Animal Welfare Ethical Review Body and the UK Home Office under the Animal (Scientific Procedures) Act 1986.

*In vivo* electrophysiological recordings were performed in 5–6 dpf zebrafish larvae from AB and Tüpfel long fin (TL) strains in accordance with the European Communities Council Directive (2010/63/EU) and French law (87/848) and approved by the Institut du Cerveau et de la Moelle épinière, the French ministry of Research and the Darwin Ethics Committee (APAFIS protocol #16469-2018071217081175v5).

### DNA cloning and transgenesis

To generate the *UAS:opsin-tdTomato* DNA constructs used for transient opsin expression and for creating the stable *Tg(UAS:opsin-tdTomato)* transgenic lines, the coding sequences of the opsins listed below and the red fluorescent protein tdTomato (from *pAAV-Syn-Chronos-tdTomato*) were cloned in frame into a UAS Tol1 backbone (*pT1UciMP*).

The source plasmids used for cloning *UAS:opsin-tdTomato* DNA constructs were:

- ChrimsonR from *pCAG-ChrimsonR-tdT* (Addgene plasmid # 59169)
- Chronos from *pAAV-Syn-Chronos-tdTomato* (Addgene plasmid # 62726)
- CoChR from *pAAV-Syn-CoChR-GFP* (Addgene plasmid # 59070)
- CheRiff from *FCK-CheRiff-eGFP* (Addgene plasmid # 51693)
- GtACR1 from *pFUGW-hGtACR1-EYFP* (Addgene plasmid # 67795)
- GtACR2 from *pFUGW-hGtACR2-EYFP* (Addgene plasmid # 67877)
- ChR2_(H134R)_ from *pAAV-Syn-ChR2(H134R)-GFP* (Addgene plasmid # 58880)

The *pCAG-ChrimsonR-tdT*, *pAAV-Syn-Chronos-tdTomato*, *pAAV-Syn-CoChR-GFP and pAAV-Syn-ChR2(H134R)-GFP* plasmids were gifts from Edward Boyden (Boyden *et al*., 2005; Klapoetke *et al*., 2014). The *FCK-CheRiff-eGFP* plasmid was a gift from Adam Cohen (Hochbaum *et al*., 2014). The *pFUGW-hGtACR1-EYFP* and *pFUGW-hGtACR2-EYFP* plasmids were gifts from John Spudich (Govorunova *et al*., 2015). The *pT1UciMP* plasmid was a gift from Harold Burgess (Addgene plasmid # 62215) (Horstick *et al*., 2015).

The cloning was achieved using the In-Fusion HD Cloning Plus CE kit (Clontech) with the following primers:

- ChrimsonR_fw, CTCAGCGTAAAGCCACCATGGGCGGAGCT
- Chronos_fw, CGTAAAGCCACCATGGAAACAGCC
- CoChR_fw, CTCAGCGTAAAGCCACCATGCTGGGAAACG
- CoChR_rev, TACTACCGGTGCCGCCACTGT
- CoChR_tdT_fw, ACAGTGGCGGCACCGGTAGTA
- CheRiff_fw, CTCAGCGTAAAGCCACCATGGGCGGAGCT
- CheRiff_rev, CTACCGGTGCCGCCACTTTATCTTCCTCTGTCACG
- CheRiff_tdT_fw, TAAAGTGGCGGCACCGGTAGTAGCAGTGAG
- GtACR1_fw, CTCAGCGTAAAGCCACCATGAGCAGCATCACCTGTGATC
- GtACR1_rev, CTACCGGTGCCGCGGTCTCGCCGGCTCTGG
- GtACR1_tdT_fw, CGAGACCGCGGCACCGGTAGTAGCAGTGAG
- GtACR2_fw, CTCAGCGTAAAGCCACCATGGCCTCCCAGGTCGT
- GtACR2_rev, CTACCGGTGCCGCCCTGCCGAACATTCTG
- GtACR2_tdT_fw, CGGCAGGGCGGCACCGGTAGTAGCAGTGAG
- ChR2(H134R)_fw, CTCAGCGTAAAGCCACCATGGACTATGGCGGCG
- ChR2(H134R)_rev, TACTCACTGCTACTACCGGTGCCGCCAC
- ChR2(H134R)_tdT_fw, ACCGGTAGTAGCAGTGAGTAAGG
- tdT_rev_40bp, CTCGAGATCTCCATGTTTACTTATACAGCTCATCCATGCC
- tdT_rev_45bp, CTAGTCTCGAGATCTCCATGTTTACTTATACAGCTCATCCATGCC

To generate the stable *Tg(UAS:opsin-tdTomato)* lines, purified *UAS:opsin-tdTomato* DNA constructs were first sequenced to confirm gene insertion and integrity and, subsequently, co-injected (35 ng/µl) with Tol1 transposase mRNA (80 ng/µl) into *Tg(KalTA4u508)* zebrafish embryos (Antinucci *et al*., 2019) at the early one-cell stage. Transient expression, visible as tdTomato fluorescence, was used to select injected embryos that were then raised to adulthood. Zebrafish codon-optimised *Tol1* transposase mRNA was prepared by *in vitro* transcription from NotI-linearised *pCS2-Tol1.zf1* plasmid using the SP6 transcription mMessage mMachine kit (Life Technologies). The *pCS2-Tol1.zf1* was a gift from Harold Burgess (Addgene plasmid # 61388) (Horstick *et al*., 2015). RNA was purified using the RNeasy MinElute Cleanup kit (Qiagen). Germ line transmission was identified by mating sexually mature adult fish to *mitfa^w2/w2^* fish and subsequently examining their progeny for tdTomato fluorescence. Positive embryos from a single fish were then raised to adulthood. Once this second generation of fish reached adulthood, positive embryos from a single ‘founder’ fish were again selected and raised to adulthood to establish stable *Tg(KalTA4u508;UAS:opsin-tdTomato)* double-transgenic lines.

To generate the *UAS:opsin-eYFP* DNA constructs used for creating the stable *Tg(UAS:opsin-eYFP)* transgenic lines, the coding sequences of the opsins fused with eYFP listed below were cloned into a UAS Tol2 backbone (*pTol2 14xUAS:MCS*). *eArch3.0-eYFP* from *pAAV-CaMKIIa-eArch_3.0-EYFP*

- *eNpHR3.0-eYFP* from *pAAV-Ef1a-DIO-eNpHR 3.0-EYFP*
- The *pAAV-CaMKIIa-eArch_3.0-EYFP* and *pAAV-Ef1a-DIO-eNpHR 3.0-EYFP* plasmids were gifts from Karl Deisseroth (Gradinaru *et al*., 2010; Mattis *et al*., 2011).

The coding sequences were amplified by PCR using the following primers and cloned into either EcoRI/NcoI (for eArch3.0) or EcoRI/SphI (for eNpHR3.0) sites of the *pTol2 14xUAS:MCS* plasmid:

- eArch3.0_fw, ATGAATTCGCCACCATGGACCCCATCGCTCT
- eArch3.0_rev, ATGCATGCTCATTACACCTCGTTCTCGTAG
- eNpHR3.0_fw, ATGAATTCGCCACCATGACAGAGACCCTGC
- eNpHR3.0_rev, TACCATGGTTACACCTCGTTCTCGTAGC

To generate the stable *Tg(UAS:opsin-eYFP)* lines, purified *UAS:opsin-eYFP* DNA constructs were first sequenced to confirm gene insertion and integrity and, subsequently, co-injected (25 ng/µl) with Tol2 transposase mRNA (25 ng/µl) into *Tg(isl2b:GAL4-VP16, myl7:TagRFP)zc65* (Fujimoto *et al*., 2011) (for eArch3.0-eYFP) or *Tg(s1020t:GAL4)* (Scott et al., 2007) (for eNpHR3.0-eYFP) zebrafish embryos at the early one-cell stage. Transient expression, visible as eYFP fluorescence, was used to select injected embryos that were then raised to adulthood. Zebrafish codon-optimised *Tol2* transposase mRNA was prepared by *in vitro* transcription from NotI-linearised *pCS2-zT2TP* plasmid using the SP6 transcription mMessage mMachine kit (Life Technologies). The *pCS2-zT2TP* was a gift from Koichi Kawakami (Suster *et al*., 2011). RNA was purified using the NucleoSpin Gel and PCR Clean-up kit (Macherey-Nagel). Germ line transmission was identified by mating sexually mature adult fish to *mitfa^w2/w2^* fish and, subsequently, examining their progeny for eYFP fluorescence. Positive embryos from each injected fish were then raised to adulthood. Once this second generation of fish reached adulthood, positive embryos from a single ‘founder’ fish were again selected and raised to adulthood to establish stable *Tg(Isl2b:GAL4;UAS:eArch3.0-eYFP) or Tg(s1020t:GAL4;UAS:eNpHR3.0-eYFP)* double-transgenic lines.

### Fluorescence image acquisition

Zebrafish embryos or larvae were mounted in 1% low-melting point agarose (Sigma-Aldrich) and anesthetised using tricaine (MS-222, Sigma-Aldrich). Imaging was performed using a custom-built 2-photon microscope [XLUMPLFLN 20× 1.0 NA objective (Olympus), 580 nm PMT dichroic, band-pass filters: 510/84 (green), 641/75 (red) (Semrock), R10699 PMT (Hammamatsu Photonics), Chameleon II ultrafast laser (Coherent Inc)]. Imaging was performed at 1040 nm for opsin-tdTomato lines, while 920 nm excitation was used for opsin-eYFP lines. In both cases, the same laser power at sample (10.7 mW) and PMT gain were used. For the images displayed in Figure 1C, 3B and 7B and Figure 7– figure supplement 3B, equivalent imaging field of view and pixel size were used (1200 × 800 px, #m/px). The imaging field of view and pixel size for images displayed in Figure 2C and 6B were 960 × 680 px, 0.385 #m/px. For all these images, the same acquisition averaging (mean image from 12 frames) and z-spacing of imaging planes (2 *μ*m) were used.

The image displayed in Figure 4A was acquired from a single plane on a fluorescence microscope [AxioExaminer D1 (Zeiss), 63× 1.0 NA objective (Zeiss), Xcite (Xcelitas, XT600) 480 nm LED illumination, 38HE filtercube (Zeiss), ImagEM camera (Hammamatsu)], with an imaging field of view of 512 × 512 px and 0.135 #m/px pixel size.

### Behavioural assays

The same monitoring system was used for all behavioural assays (see schematic in Figure 2A) with some differences. Images were acquired under infrared illumination (850 nm) using a high-speed camera (Mikrotron MC1362, 500 µs shutter-time) equipped with a machine vision lens (Fujinon HF35SA-1) and an 850 nm bandpass filter to block visible light. The 850 nm bandpass filter was removed during embryonic activation assays (in which images were acquired at 1,000 fps) to determine time of light stimulus onset. In all other assays, lower acquisition rates were used (i.e. 50 or 500 fps) and, within each assay, the frames corresponding to stimulus onset/offset were consistent across trials.

Light was delivered across the whole arena from above using the following LEDs:

*For embryonic assays*

- 470 nm OSRAM Golden Dragon Plus LED (LB W5AM).
- 590 nm ProLight LED (PM2B-3LAE-SD).

*For larval assays*

- 459 nm OSRAM OSTAR Projection Power LED (LE B P2W).
- 617 nm OSRAM OSTAR Projection Power LED (LE A P2W).

The 459 and 617 nm LEDs were projected onto the arena with an aspheric condenser with diffuser surface. Irradiance was varied using constant current drive electronics with pulse-width modulation at 5 kHz. Irradiance was calibrated using a photodiode power sensor (Thorlabs S121C). LED and camera control were implemented using LabVIEW (National Instruments).

Before experiments, animals were screened for opsin expression in the target neural population at either 22 hpf (embryonic assays) or 3 dpf (larval assays) using a fluorescence stereomicroscope (Olympus MVX10). For each opsin, animals with similar expression level were selected for experiments together with control opsin-negative siblings. To reduce variability in opsin expression level, all animals used for behavioural experiments were heterozygous for both the GAL4 and UAS transgenes. Animals were placed in the arena in the dark for around 2 min before starting experiments. For all assays, each light stimulus was repeated at least 3 times. Each trial lasted 1 s in behavioural activation assays and 30 s in behavioural inhibition assays.

### Embryonic activation assay

Opsin expression was targeted to trigeminal ganglion neurons using the *Tg(isl2b:GAL4)* transgene (Ben Fredj *et al*., 2010). Behaviour was monitored at 1,000 fps across embryos (28–30 hpf) individually positioned in agarose wells (∼2 mm diameter) in fish facility water and free to move within their chorion. Embryos were subjected to 5 or 40 ms pulses of blue (470 nm) or amber (590 nm) light at different irradiance levels (4.5–445 *μ*W/mm^2^) and with a 15 s inter-stimulus interval in the dark.

### Embryonic inhibition assay

Opsin expression was targeted to spinal primary and secondary motor neurons and interneurons (Kolmer-Agduhr cells and ventral longitudinal descending interneurons) using the *Tg(s1020t:GAL4)* transgene (Scott *et al*., 2007). Behaviour was monitored at 50 fps across embryos (24–27 hpf) individually positioned in agarose wells (∼2 mm diameter) with fish facility water and free to move within their chorion. Embryos were subjected to 10 s pulses of blue (470 nm) or amber (590 nm) light at different irradiance levels (0–227 *μ*W/mm^2^) with a 50 s inter-stimulus interval in the dark.

### Larval activation assay

Opsin expression was targeted to primary and secondary spinal motor neurons using the *Tg(mnx1:GAL4)* transgene (Bohm *et al*., 2016). Behaviour was monitored at 500 fps in 6 dpf larvae with their head restrained in 2% low-melting point agarose (Sigma-Aldrich) and their tail free to move. Larvae were subjected to 2 or 10 ms pulses of blue (459 nm) or red (617 nm) light at different irradiance levels (0.04–2.55 mW/mm^2^) with a 20 s inter-stimulus interval in the dark. We also provided 250 ms trains of light pulses (1 ms pulse duration for blue light at 2.55 mW/mm^2^ or 10 ms for red light at 1 mW/mm^2^) at two pulse frequencies (20 or 40 Hz).

### Larval inhibition assays

Opsin expression was targeted to spinal cord neurons using either the *Tg(s1020t:GAL4)* or *Tg(mnx1:GAL4)* transgene, as above. Behaviour was monitored at 50 fps across larvae individually positioned in agarose wells (∼1.4 cm diameter) with fish facility water in which they were free to swim. Larvae were subjected to 10 s pulses of blue (459 nm) or red (617 nm) light at different irradiance levels (0.24–2.55 mW/mm^2^) with a 50 s inter-stimulus interval in the dark. Control trials during which no light pulse was provided were interleaved between light stimulation trials.

### Behavioural data analysis

Movie data was analysed using MATLAB (MathWorks). Region of interests (ROIs) containing individual fish were manually specified. For each ROI, the frame-by-frame change in pixel intensity – ΔPixel – was computed in the following way. For each trial, pixel intensity values were low-pass filtered across time frames and the absolute frame-by-frame difference in intensity (*dI*) was obtained for each pixel. Pixels showing the highest variance in *dI* (top 5^th^ percentile) were selected to compute their mean *dI,* corresponding to the ROI ΔPixel trace for the trial.

With the exception of the larval inhibition assay (see below), onset and offset of animal movements were detected from ΔPixel traces in the following way. For each ROI, ΔPixel traces were concatenated across all trials to estimate the probability density function (*pdf*) of ΔPixel values. The portion of the distribution with values below the *pdf* peak was mirror-reflected about the *x*-axis and a Gaussian was fitted to the obtained symmetric distribution. The mean (*μ*) and standard deviation (σ) of the fitted Gaussian were then used to compute ROI-specific ΔPixel thresholds for detecting onset (*μ* + 6σ) and offset (*μ* + 3σ) of animal movements.

For embryonic and larval activation assays, behavioural response latency corresponds to the time from light stimulus onset to the start of the first detected movement. Movements were classified as optogenetically-evoked if their response latency was shorter than 200 ms for the embryonic assay or 50 ms for the larval assay, which corresponds to the minimum in the *pdf* of response latency from all opsin-expressing larvae (Figure 3E). For each animal, response probability to each light stimulus type corresponds to the fraction of trials in which at least one optogenetically-evoked movement was detected.

In the larval activation assay, the tail was tracked by performing consecutive annular line-scans, starting from a manually-selected body centroid and progressing towards the tip of the tail so as to define nine equidistant x-y coordinates along the tail. Inter-segment angles were computed between the eight resulting segments. Reported tail curvature was computed as the sum of these inter-segment angles. Rightward bending of the tail is represented by positive angles and leftward bending by negative angles. Number of tail beats corresponds to the number of full tail oscillation cycles. Tail theta-1 angle is the amplitude of the first half beat. Tail beat frequency was computed as the reciprocal of the mean full-cycle period during the first four tail oscillation cycles of a swim bout. Bout duration was determined from ΔPixel traces using the movement onset/offset thresholds described above.

For larval inhibition assays, images were background-subtracted using a background model generated over each trial (30 s duration). Images were then thresholded and the fish body centroid was found by running a particle detection routine for binary objects within suitable area limits. Tracking of body centroid position was used to compute fish speed, and periods in which speed was higher than 1 mm/s were classified as swim bouts. Bout speed was computed as the mean speed over the duration of each bout.

To account for group differences in baseline coil/bout rate and bout speed in inhibition assays, data was normalised at a given irradiance level by divided by the mean rate/speed across fish in control (no light) trials.

### Electrophysiological recordings

#### Transgenic lines

Opsin expression was targeted to primary motor neurons using the *Tg(mnx1:GAL4)* transgene (Bohm *et al*., 2016) with one exception: 11 out of 19 eNpHR3.0-expressing cells were recorded in *Tg(s1020t:GAL4)* larvae (Scott *et al*., 2007). As in behavioural assays, all animals used for electrophysiological experiments were heterozygous for both the GAL4 and UAS transgenes. For control recordings, we targeted opsin-negative GFP-expressing primary motor neurons in *Tg(mnx1:GAL4;UAS:EGFP)* (Asakawa *et al*., 2008) or *Tg(parga-GFP)* (Balciunas et al., 2004) larvae. In all transgenic lines used, primary motor neurons could be unambiguously identified as the 3–4 largest cell somas, located in the dorsal-most portion of the motor column (Beattie *et al*., 1997; Bello-Rojas *et al*., 2019). We verified primary motor neuron identity in a small subset of recordings from eYFP-expressing cells in *Tg(mnx1:GAL4;UAS:ChR2(H134R)-eYFP)* larvae by adding 0.025% sulforhodamine-B acid chloride dye in the intracellular solution (Sigma-Aldrich) and filling the neuron to reveal its morphology. To maximise data acquisition in an in vivo preparation, when the first attempts of primary motor neuron recordings were not successful, we recorded neighbouring, dorsal located presumed secondary motor neurons (11 out of 86 included cells).

#### Data acquisition

Zebrafish larvae (5–6 dpf) were first paralysed in 1 mM α-Bungarotoxin solution (Tocris) for 3–6 min after which they were pinned in a lateral position to a Sylgard-coated recording dish (Sylgard 184, Dow Corning) with tungsten pins inserted through the notochord. The skin was removed between the trunk and midbody regions using sharp forceps, after which the dorsal muscle from 2–3 somites was suctioned with glass pipettes (∼50 µm opening made from capillaries of 1.5 mm outer diameter, 1.1 mm inner diameter; Sutter). Patch pipettes were made from capillary glass (1 mm outer diameter, 0.58 mm inner diameter; WPI) with a horizontal puller (Sutter Instrument P1000) and had resistances between 8–16 M&. To first pass the dura, we applied a higher positive pressure (30–40 mm Hg) to the recording electrode via a pneumatic transducer (Fluke Biomedical, DPM1B), which was then lowered (20–25 mm Hg) once the electrode was near the cells. We generally recorded data from a single cell per larva. In a few instances, two cells from separate adjacent somites were recorded in the same fish.

External bath recording solution contained the following: 134 mM NaCl, 2.9 mM KCl, 2.1 mM CaCl2-H_2_O, 1.2 mM MgCl_2_, 10 mM glucose, and 10 mM HEPES, with pH adjusted to 7.8 with 9 mM NaOH and an osmolarity of 295 mOsm. We blocked glutamatergic and GABAergic synaptic transmission with a cocktail of: 20 µM CNQX or DNQX, 50 µM D-AP5, 10 µM Gabazine (Tocris) added to the external recording solution. The –50 mV ECl solution contained: 115 mM K-gluconate, 15 mM KCl, 2 mM MgCl_2_, 4 mM Mg-ATP, 0.5 mM EGTA, 10 mM HEPES, with pH adjusted to 7.2 with 11mM KOH solution, and a 285 mOsm. In these conditions, we calculated the liquid junction potential (LJP; Clampfit calculator) to be 12.4 mV. The –70 mV ECl solution contained: 126 mM K-gluconate, 4 mM KCl, 2 mM MgCl_2_, 4 mM Mg-ATP, 0.5 mM EGTA, 10 mM HEPES, pH adjusted to 7.2 with 11mM KOH solution, 285 mOsm and a 13.3 mV LJP. All reagents were obtained from Sigma-Aldrich unless otherwise stated.

Recordings were made with an Axopatch 700B amplifier and digitised with Digidata 1440A or 1550B (Molecular Devices). pClamp software was used to acquire electrophysiological data at a sampling rate of 20 kHz and low-pass filtered at 2 kHz (voltage clamp) or 10 kHz (current clamp). Voltage clamp recordings were acquired with full whole-cell compensation and ∼60% series resistance compensation, while corrections for bridge balance and electrode capacitance were applied in current clamp mode. Cells were visualised with a 63×/1.0 NA or a 60×/1.0 NA water-immersion objective (Zeiss or Nikon, respectively) on a fluorescence microscope equipped with differential interference contrast optics (AxioExaminer D1, Zeiss or Eclipse FN1, Nikon).

#### Optogenetic stimulation

Light stimulation was performed with either a X-Cite (Xcelitas, XT600) or a broadband white LED (Prizmatix, UHP-T-HCRI_DI) light source equipped with a combination of different bandpass and neutral density filters to modulate irradiance at specific wavelengths (see Figure 4–figure supplement 1A for wavelengths and irradiance levels used to activate opsins). The onset, duration and irradiance level of light pulses were triggered and controlled via the Digidata device used for electrophysiological recordings.

For all cells, data was acquired in the following order: (1) series resistance was checked at the beginning, middle and end of recording; (2) action potential rheobase was determined by injecting 5 ms pulses of current (160–340 pA) in current-clamp gap-free mode; (3) voltage clamp recording of opsin photocurrents; (4) current clamp recording of voltage responses induced by opsin activation. Light stimuli were provided from low to high irradiance levels across all protocols. For each protocol, inter-stimulus intervals were between 10 and 15 s.

For cation channelrhodopsins, we used a range of short light pulses. Voltage clamp recordings were paired with a 5 ms light pulse, while current clamp recordings were performed with 1, 2 or 5 ms-long pulses. In addition, we tested whether we could optogenetically entrain neurons to spike at frequencies ranging from 1 to 100 Hz using stimulus trains composed of 2 or 5 ms-long light pulses.

For anion channelrhodopsins and Cl^−^/H^+^ pumps, voltage and current clamp recordings were paired with a 1 s light pulse. In addition, we used two different tests of optogenetic inhibition during active spiking. To assess single spike inhibition efficacy and precision, we evoked spiking by injecting 5 ms pulses of current at 1.2–1.5× rheobase for 10 trains at 5 Hz (1 s inter-train interval, total of 100 spikes triggered in 30 s), during which we provided 5 ms-long light pulses paired to the first current stimulus of the train and a subsequent one with progressively longer latency (Zhang *et al*., 2007). To test opsin ability to inhibit tonic firing over longer time periods, we evoked spiking with longer pulses of current (200–800 ms) at 1.2–1.5x rheobase paired with a light pulse (50–200 ms-long) in the middle of the current stimulation. We first recorded a control current injection-only trial, followed by current and light pulse trials with a 20 s inter-stimulus interval.

## Data analysis

Data were analysed using the pyABF module in Spyder (3.3.6 MIT, running Python 3.6, scripts available here: https://github.com/wyartlab/Antinucci_Dumitrescu_et_al_2020), MATLAB (MathWorks) and Clampfit (Molecular Devices). Series resistance (Rs) was calculated as a cell response to a 5 or 10 mV hyperpolarisation step in voltage clamp from a holding potential of – 60 mV, with whole-cell compensation disabled. Membrane resistance (Rm) was obtained from the steady holding current at the new step, and membrane capacitance (Cm) corresponds to the area under the exponentially decaying current from peak to holding. We used the following cell inclusion criteria:

(1) cell spiking upon injection of a 5 ms-long pulse of current; (2) membrane resting potential < – 50 mV at all times; (3) > 150 pA current injection necessary to maintain the cell at a holding potential equal to resting potential in current clamp; (4) series resistance < 6× pipette resistance at all times during the recording. We chose this conservative series resistance range as per previous electrophysiological procedures in other animal models: i.e. mammalian *in vivo* recordings with pipette resistance between 4–7 MΩ and max series resistance between 10–100 MΩ (Margrie *et al*., 2002). All reported membrane voltages were liquid junction potential corrected.

For voltage clamp recordings, we measured the maximum photocurrent amplitude in a time window of 100 ms (for cation channelrhodopsins) or 1 s (for anion channelrhodopsins and Cl^−^/H^+^ pumps) duration starting from light onset. To characterise photocurrent kinetics of cation channelrhodopsins, we measured the time to peak photocurrent from light onset (i.e. activation time) and computed the response decay time constant by fitting a monoexponential decay function to the photocurrent from peak to baseline (i.e. deactivation time constant). To compute photocurrent kinetics of anion channelrhodopsins and Cl^−^/H^+^ pumps, we fitted monoexponential functions to the following components of the response: activation time constant was computed from light onset to peak response, inactivation time constant from peak response to steady state (last 5 ms of light stimulation), deactivation time constant from steady state to baseline (1 s following light offset)

To characterise voltage responses induced by opsins under current clamp, we first classified events as spikes (when max voltage depolarisation was > – 30 mV) or sub-threshold (peak voltage deflection < – 30 mV). For each response type, we measured the absolute peak of the response, the time to reach maximum response from light onset and the time-decay to baseline from peak by fitting a monoexponential decay function, as above. To assess firing pattern fidelity, we calculated the number of spikes per light pulse in a train, the latency from light onset to the first spike occurring within a 10 ms time window, and the spike jitter as the standard deviation of spike latency values across a pulse train with given frequency.

Opsin efficacy in inhibiting single spikes was quantified using the following equation:

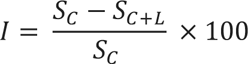

where *S_C_* is the mean number of spikes elicited by current pulses when no light was provided, *SC+L* is the mean number of spikes elicited during time periods in which a light pulse was paired with a current pulse, and *I* is the inhibition index (100% being perfect inhibition and negative values indicating additional spikes were generated during light pulses). Tonic firing inhibition efficacy was quantified by counting the number of spikes occurring during the light delivery period and normalising this count to provide spikes generated per 50 ms.

### Statistical analysis

All statistical analyses were performed using Prism (GraphPad). Sample distributions were first assessed for normality and homoscedasticity. Details regarding the statistical tests used are reported in Supplementary File 2 for behavioural data and Supplementary File 3 for electrophysiological data. Significance threshold was set to 0.05 and all reported p-values were corrected for multiple comparisons. Tests were two-tailed for all experiments. Number of animals/cells are provided for each graph. No outliers were excluded from the analyses.

**Video 1. Escape responses elicited by optogenetic stimulation of embryonic trigeminal neurons** Escape responses in *Tg(isl2b:GAL4;UAS:CoChR-tdTomato)* embryos (28–30 hpf) triggered by a 5 ms pulse of blue light (470 nm, 445 *μ*W/mm^2^). Images were acquired at 1,000 frames per second and the video plays at 0.1× speed. Related to Figure 2.

Figure 2–Source Data 1. Data related to Figure 2.

Data provided as a XLSX file.

**Video 2. Swim bouts elicited by single-pulse optogenetic stimulation of larval spinal motor neurons**

Swim responses in 3 head-restrained tail-free *Tg(mnx1:GAL4;UAS:CoChR-tdTomato)* larvae (6 dpf, left) triggered by a single 2 ms pulse of blue light (459 nm, 0.63 mW/mm^2^). A control opsin-negative larva is positioned on the right. Images were acquired at 500 frames per second and the video plays at 0.04× speed. Related to Figure 3.

**Video 3. Swim bouts elicited by 20 Hz pulse train optogenetic stimulation of larval spinal motor neurons**

Swim responses in 3 head-restrained tail-free *Tg(mnx1:GAL4;UAS:CoChR-tdTomato)* larvae (6 dpf, left) triggered by a train of 1 ms pulses of blue light (459 nm, 20 Hz, 2.55 mW/mm^2^, 250 ms train duration). A control opsin-negative larva is positioned on the right. Images were acquired at 500 frames per second and the video plays at 0.04× speed. Related to Figure 3.

Figure 3–Source Data 1. Data related to Figure 3.

Data provided as a XLSX file.

Figure 4–Source Data 1. Data related to Figure 4.

Data provided as a XLSX file.

Figure 5–Source Data 1. Data related to Figure 5.

Data provided as a XLSX file.

Video 4. Monitoring of coiling behaviour upon opsin activation in embryonic spinal neurons

Coiling behaviour in *Tg(s1020t:GAL4;UAS:GtACR2-tdTomato)* embryos (24–27 hpf) subjected to a 10 s period of blue light (470 nm, 225 *μ*W/mm^2^). Images were acquired at 50 frames per second and the video plays at 3× speed. Related to Figure 6.

Figure 6–Source Data 1. Data related to Figure 6.

Data provided as a XLSX file.

Video 5. Suppression of swimming upon opsin activation in larval spinal neurons

Suppression of swimming in *Tg(s1020t:GAL4;UAS:GtACR1-tdTomato)* larvae (6 dpf) during 10 s of blue light (459 nm, 0.24 mW/mm^2^). Images were acquired at 50 frames per second and the video plays at 3× speed. Related to Figure 7.

*Figure 7–Source* Data 1. Data related to Figure 7.

Data provided as a XLSX file.

Figure 8–Source Data 1. Data related to Figure 8.

Data provided as a XLSX file.

Figure 9–Source Data 1. Data related to Figure 9.

Data provided as a XLSX file.

---

**Figure 2–figure supplement 1.**
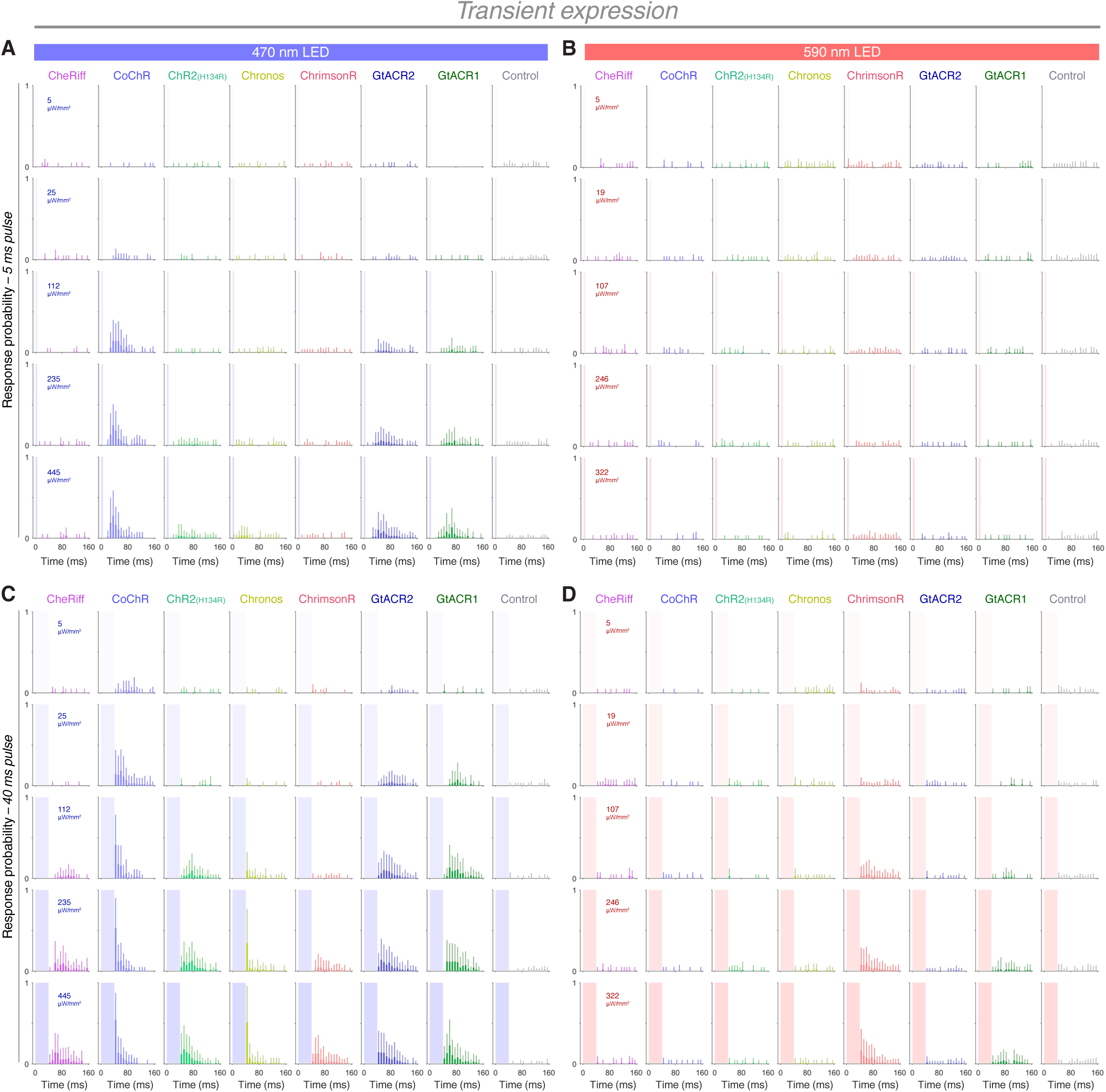
Response probability vs. time in transient transgenic embryos expressing opsins in trigeminal neurons. **A–D** Distribution of response probability vs. time for *Tg(isl2b:GAL4)* embryos (28–30 hpf) expressing different opsins through transient transgenesis (mean + SD, across fish). Embryos were stimulated with 5 ms (**A,B**) or 40 ms (**C,D**) pulses of blue (470 nm; **A,C**) or amber (590 nm; **B,D**) light. Each time bin corresponds to 8 ms.

**Figure 2–figure supplement 2.**
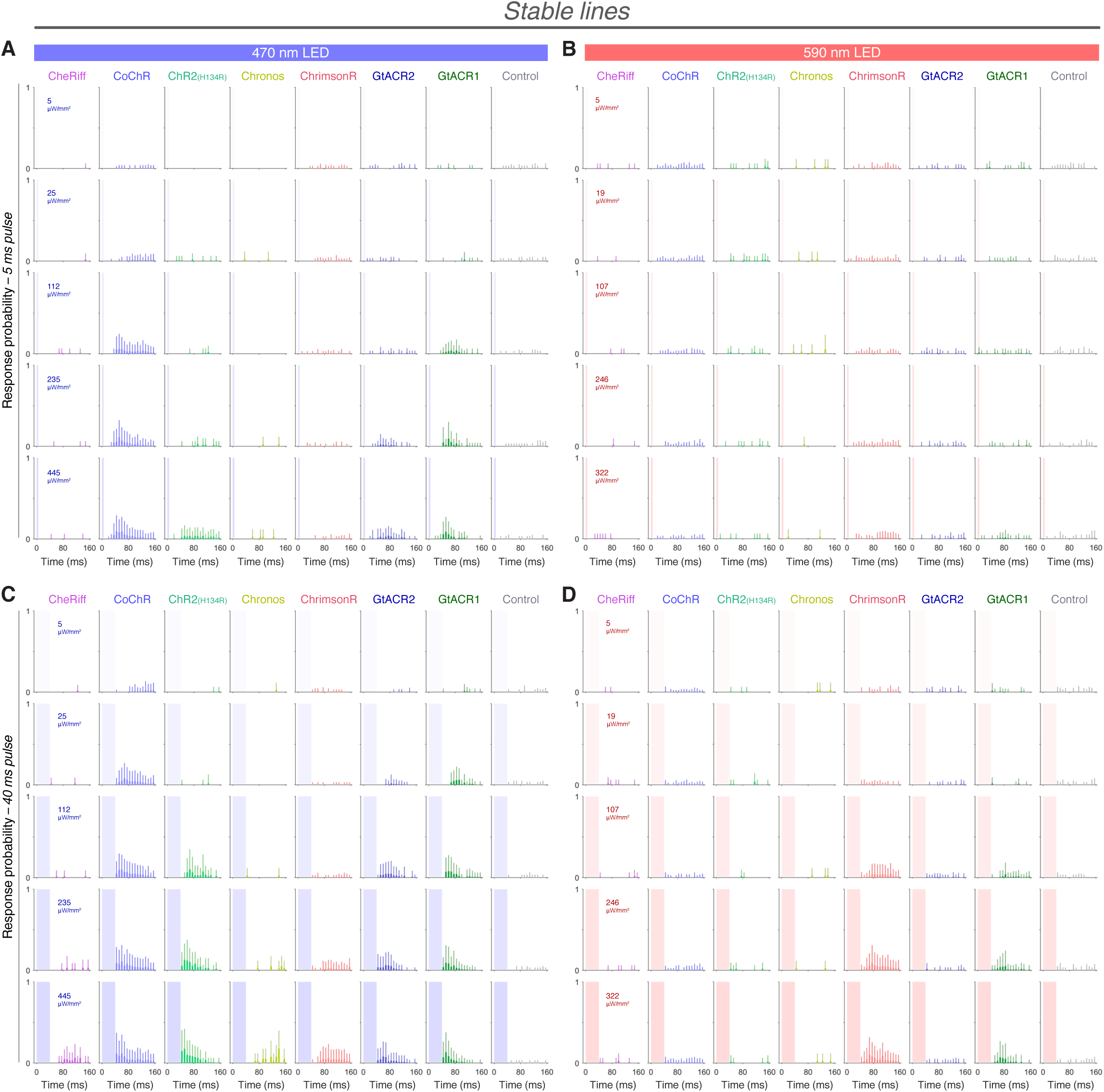
Response probability vs. time in stable transgenic embryos expressing opsins in trigeminal neurons. **A–D** Distribution of response probability vs. time for *Tg(isl2b:GAL4)* embryos (28–30 hpf) expressing different opsins through stable transgenesis (mean + SD, across fish). Embryos were stimulated with 5 ms (**A,B**) or 40 ms (**C,D**) pulses of blue (470 nm; **A,C**) or amber (590 nm; **B,D**) light. Each time bin corresponds to 8 ms.

**Figure 3–figure supplement 1.**
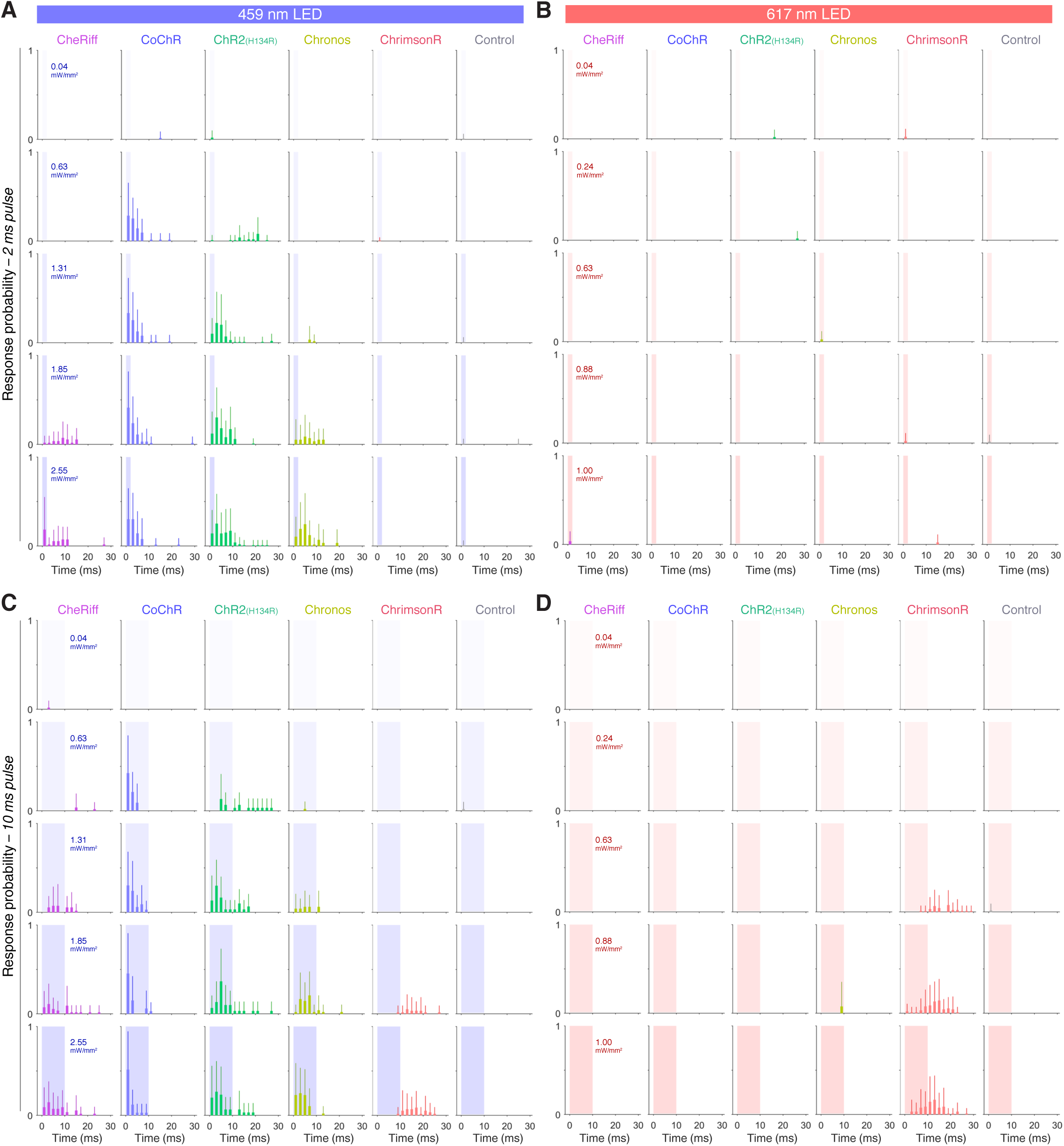
Response probability vs. time in larvae expressing opsins in spinal motor neurons. **A–D** Distribution of response probability vs. time for *Tg(mnx1:GAL4)* larvae (6 dpf) expressing different opsins (mean + SD, across fish). Larvae were stimulated with single 2 ms (**A,B**) or 10 ms (**C,D**) pulses of blue (459 nm; **A,C**) or red (617 nm; **B,D**) light. Each time bin corresponds to 2 ms.

**Figure 4–figure supplement 1.**
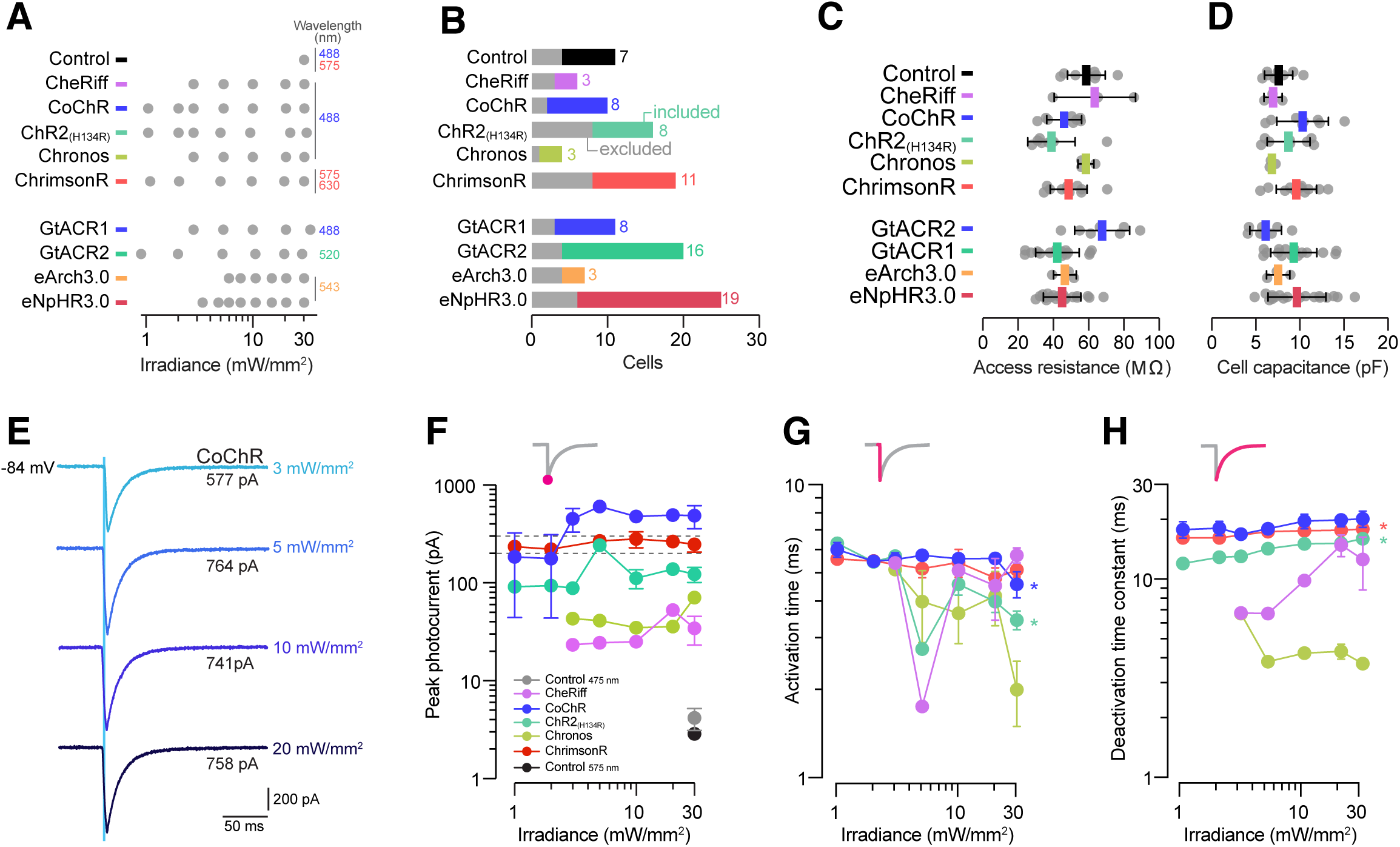
Wavelengths used in electrophysiological recordings and photocurrent properties vs. irradiance. **A** Wavelengths and irradiance levels used for each opsin line and control cells. **B** Number of cells patched in each group. Numbers and coloured bars indicate included cells while grey bars indicate excluded cells (see Materials and methods for inclusion criteria). **C,D** Access resistance (**C**) and cell capacitance (**D**) were comparable between groups (mean ± SD, across cells). **E** Example photocurrents from a CoChR-expressing cell at different irradiance levels (3– 20 mW/mm^2^). **F–H** Peak photocurrent amplitude (**F**), activation time (**G**) and deactivation time constant (**H**) vs. irradiance (mean ± SEM, across cells). Dotted lines in (**F**) show range of pMN rheobase. Asterisks indicate a significant non-zero slope.

**Figure 5–figure supplement 1.**
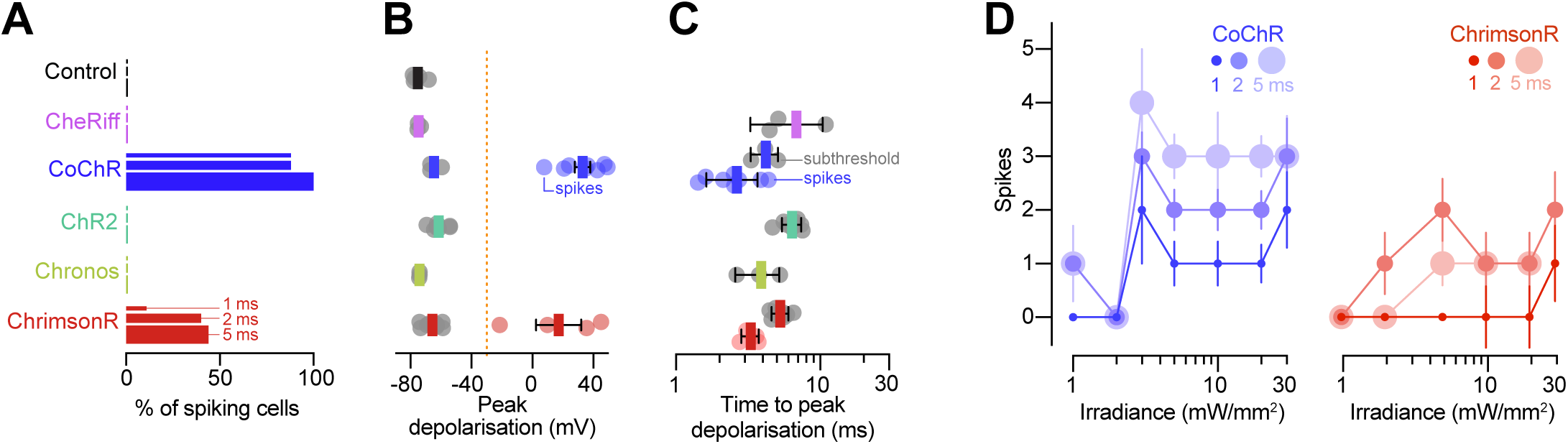
Optogenetically-evoked voltage responses. **A** Fraction of cells that generated spikes in response to single light pulses (1–5 ms). **B** Peak depolarisation across irradiance levels (1–30 mW/mm^2^; mean ± SEM, across cells). Orange line indicates threshold for spike detection (–30 mV). **C** Time to peak depolarisation (mean ± SEM). **D** Number of evoked spikes vs. irradiance (1–5 ms pulse duration). In CoChR-expressing cells, 2– 5 ms light pulses induced spike bursts (mean ± SEM).

**Figure 6–figure supplement 1.**
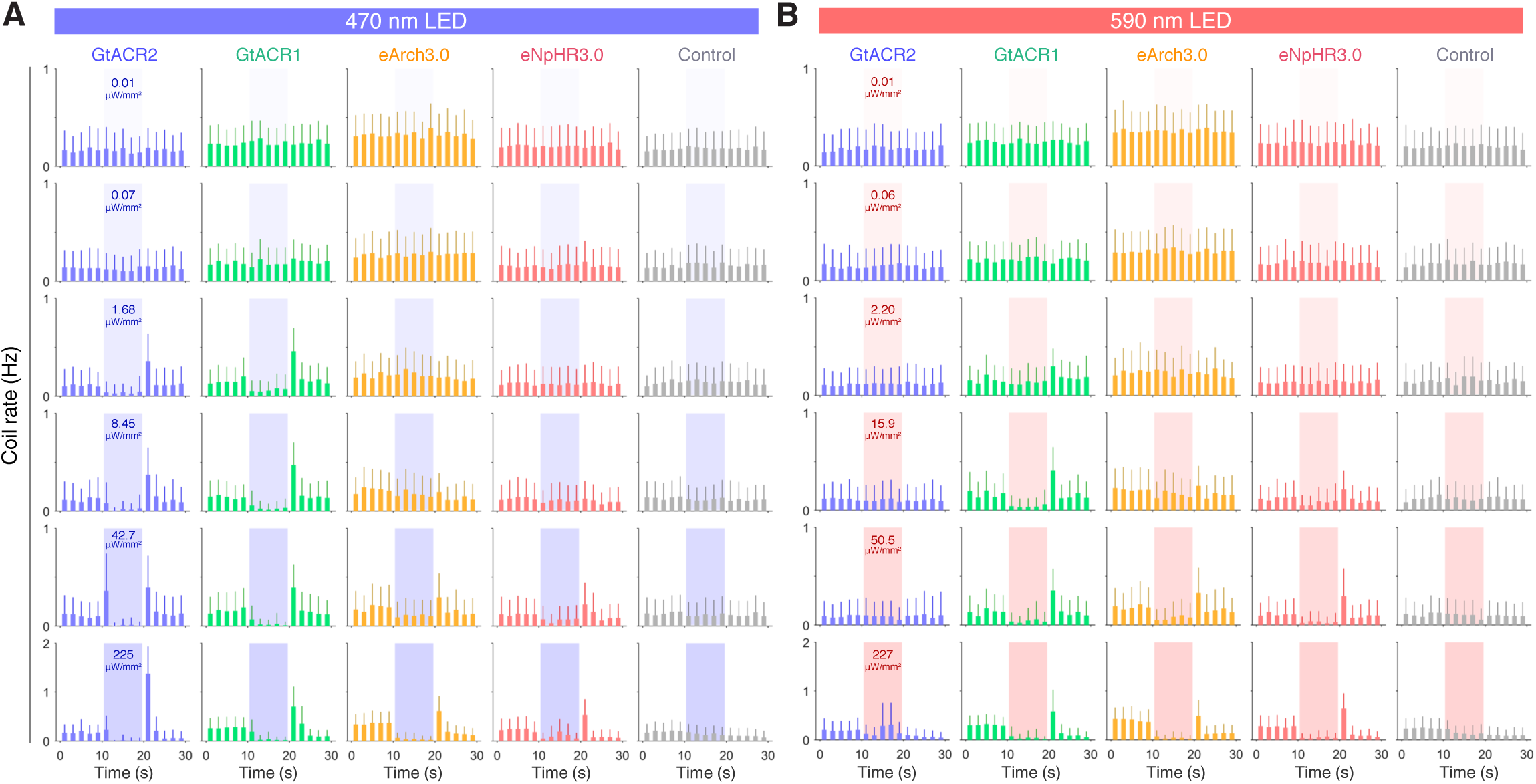
Coil rate vs. time in embryos expressing different opsins in spinal neurons. **A,B** Distribution of coil rate vs. time for *Tg(s1020t:GAL4)* embryos (24–27 hpf) expressing different opsins (mean + SD, across fish). Embryos were subjected to 10 s pulses of blue (470 nm; **A**) or amber (590 nm; **B**) light. Each time bin corresponds to 2 s.

**Figure 6–figure supplement 2.**
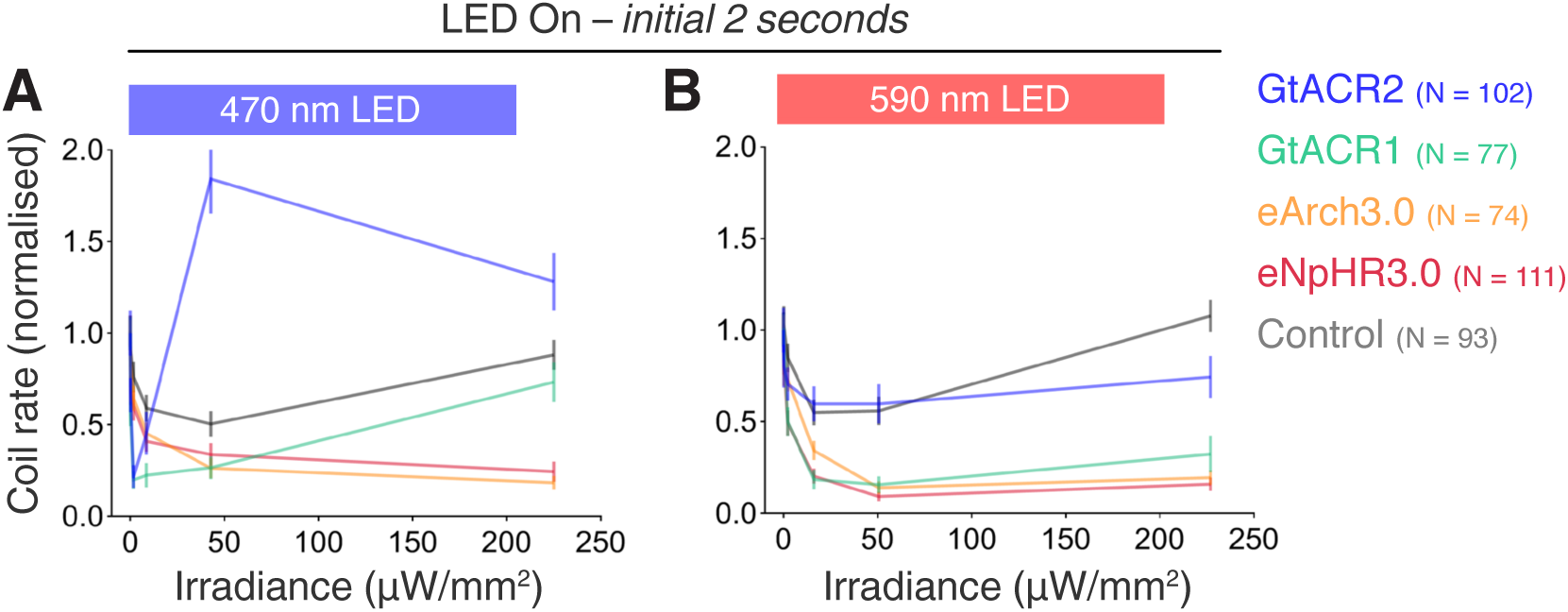
Coil rate vs. irradiance for the initial 2 seconds of light exposure. A,B Normalised coil rate during the initial 2 s of the LED On period in embryos (24–27 hpf) expressing different opsins (mean ± SEM, across fish). Control opsin-negative siblings were subjected to the same light stimuli.

**Figure 7–figure supplement 1.**
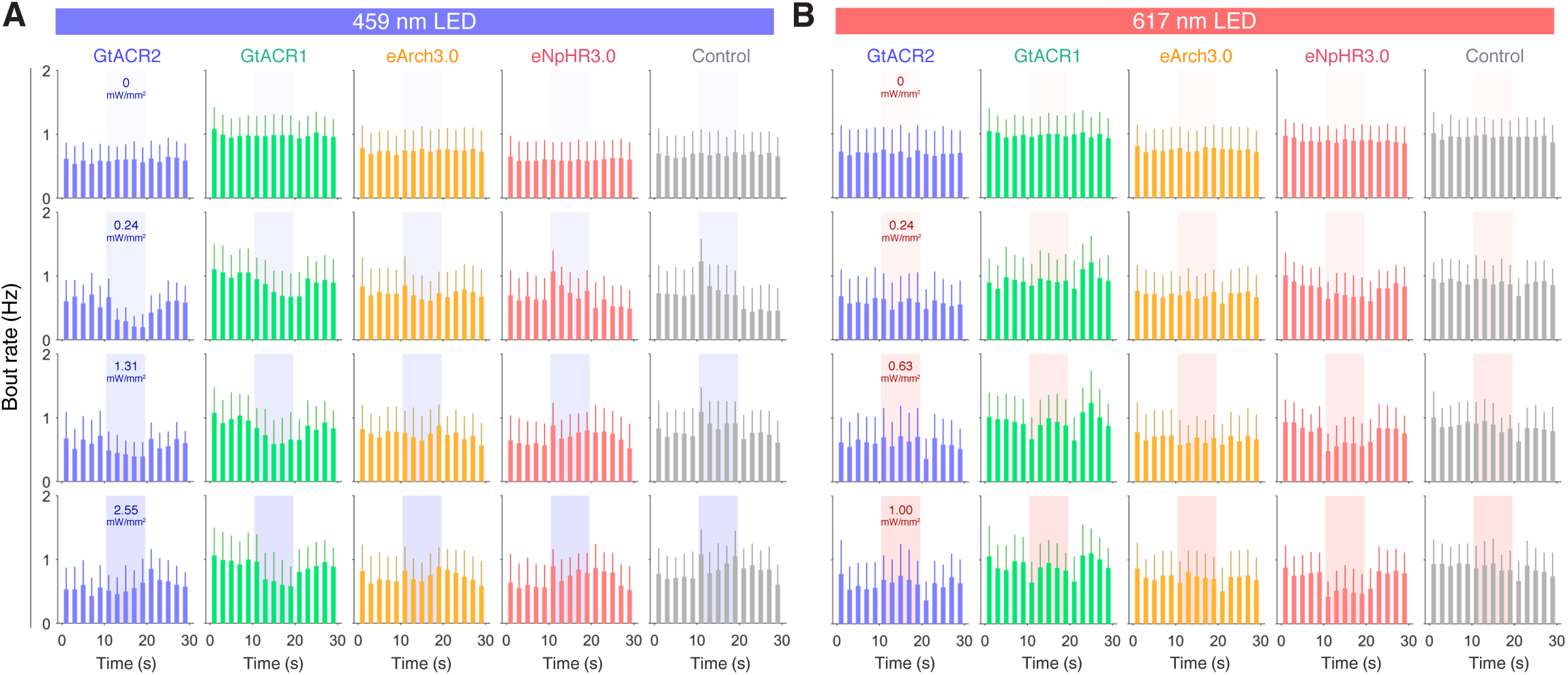
Bout rate vs. time in larvae expressing different opsins in spinal neurons. **A,B** Distribution of bout rate vs. time for *Tg(s1020t:GAL4)* larvae (6 dpf) expressing different opsins (mean + SD, across fish). Larvae were subjected to 10 s pulses of blue (459 nm; **A**) or red (617 nm; **B**) light. Each time bin corresponds to 2 s.

**Figure 7–figure supplement 2.**
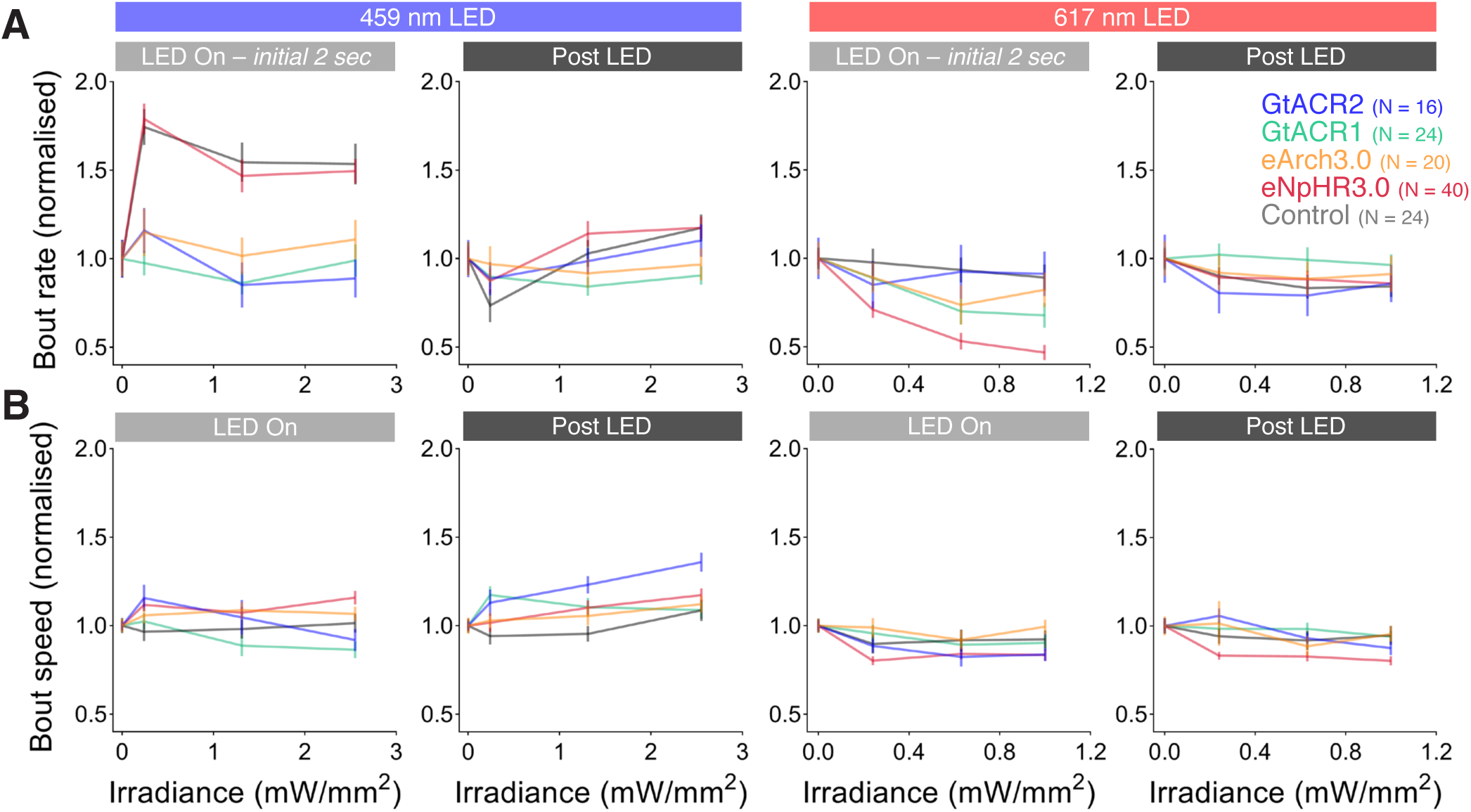
Bout rate and speed vs. irradiance during different time periods in *Tg(s1020t:GAL4)* larvae. **A,B** Normalised bout rate (**A**) or bout speed (**B**) during the whole LED On period, the initial 2 s of light exposure and the ‘post LED’ 8 s period in *Tg(s1020t:GAL4)* larvae (6 dpf) expressing different opsins (mean ± SEM, across fish). Control opsin-negative siblings were subjected to the same light stimuli.

**Figure 7–figure supplement 3.**
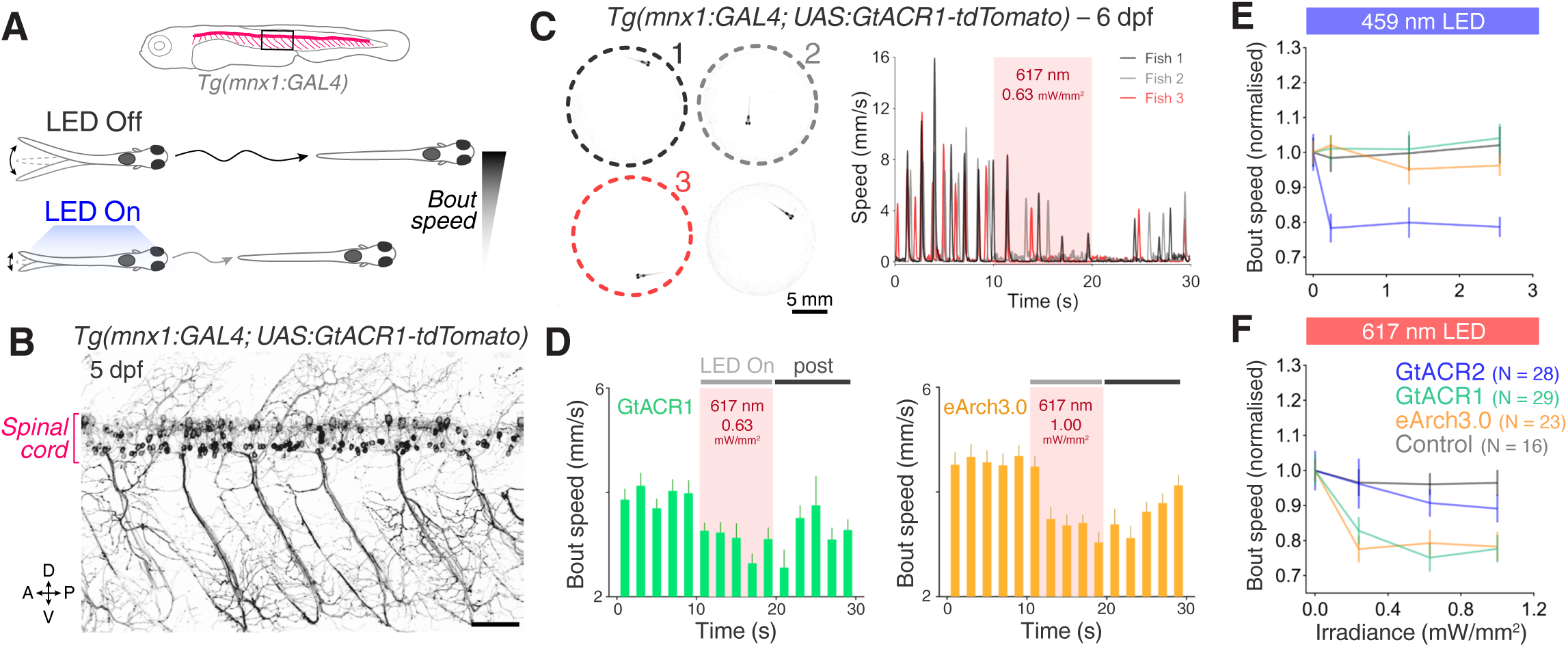
Optogenetic suppression of swimming in *Tg(mnx1:GAL4)* larvae. **A** Schematics of opsin expression pattern and behavioural assay. **B** Opsin expression in spinal motor neurons and interneurons in a *Tg(mnx1:GAL4;UAS:GtACR1-tdTomato)* larva at 5 dpf. Imaging field of view corresponds to black box in (**A**). A, anterior; D, dorsal; P, posterior; V, ventral. Scale bar 50 *μ*m. **C** Background-subtracted camera field of view showing *Tg(mnx1:GAL4;UAS:GtACR1-tdTomato)* larvae positioned in individual agarose wells (left) and tracking of swimming speed for selected larvae (right). Behaviour was monitored at 50 fps across multiple freely-swimming larvae (6 dpf; N = 24 ± 6 fish per group, mean ± SD) while they were subjected to 10 s light periods (459 or 617 nm, 0–2.55 mW/mm^2^) with a 50 s inter-stimulus interval. **D** Optogenetically-induced changes in bout rate (mean + SEM, across fish) in *Tg(mnx1:GAL4)* larvae expressing GtACR1 (N = 29 larvae, left) or eArch3.0 (N = 23 larvae, right). Horizontal grey bars indicate the time windows used for comparative quantification of behavioural changes. Each time bin corresponds to 2 s. **E,F** Normalised bout speed during the ‘LED On’ period in larvae expressing different opsins (mean ± SEM, across fish). Control opsin-negative siblings were subjected to the same light stimuli.

**Figure 7–figure supplement 4.**
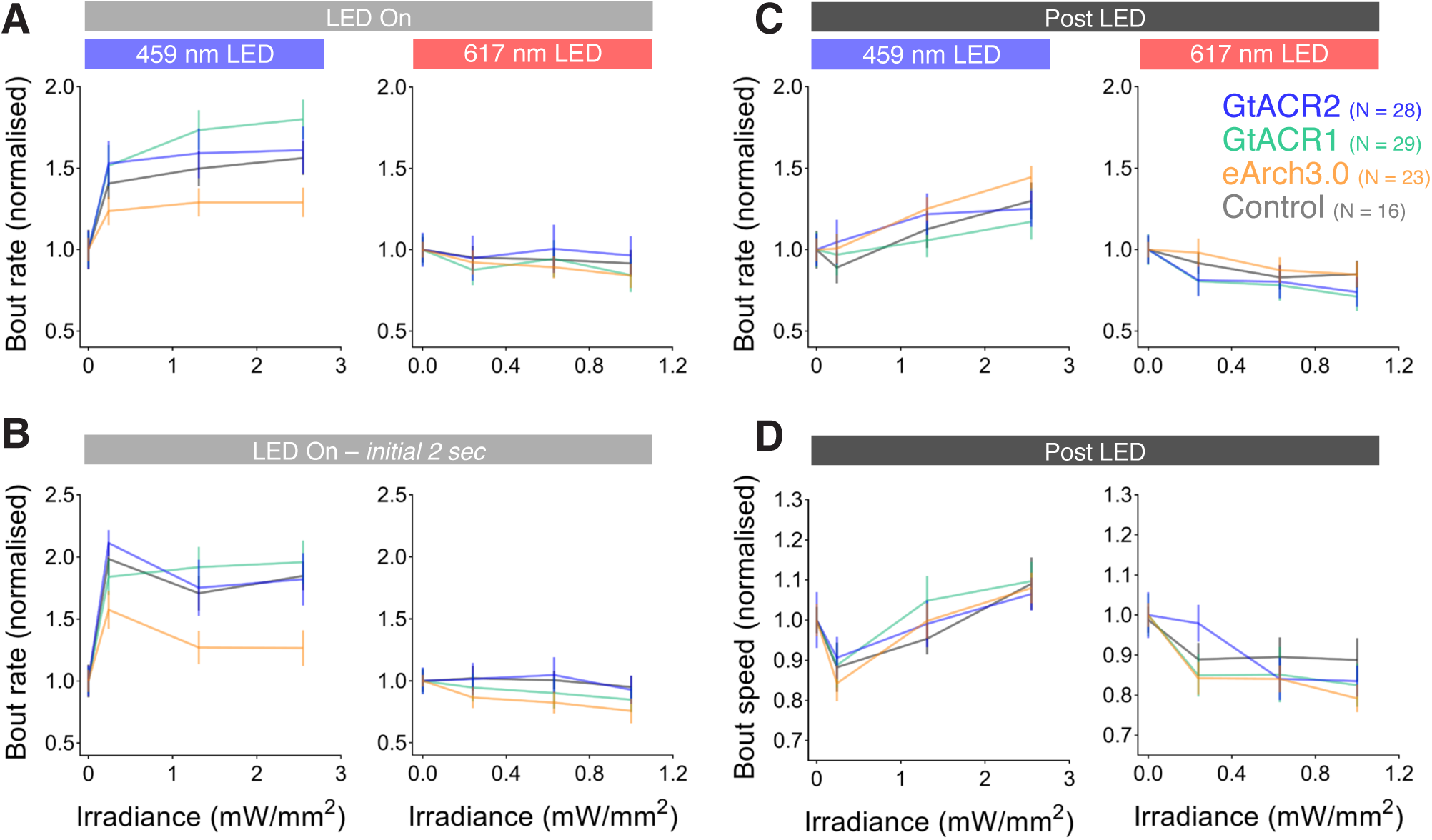
Bout rate and speed vs. irradiance during different time periods in *Tg(mnx1:GAL4)* larvae. **A–D** Normalised bout rate (**A–C**) or bout speed (**D**) during the whole ‘LED On’ period (**A**), the initial 2 s of the light period (**B**), or the ‘post LED’ 8 s period (**C,D**) in Tg(*mnx1:GAL4)* larvae (6 dpf) expressing different opsins (mean ± SEM, across fish). Control opsin-negative siblings were subjected to the same light stimuli.

**Figure 8–figure supplement 1.**
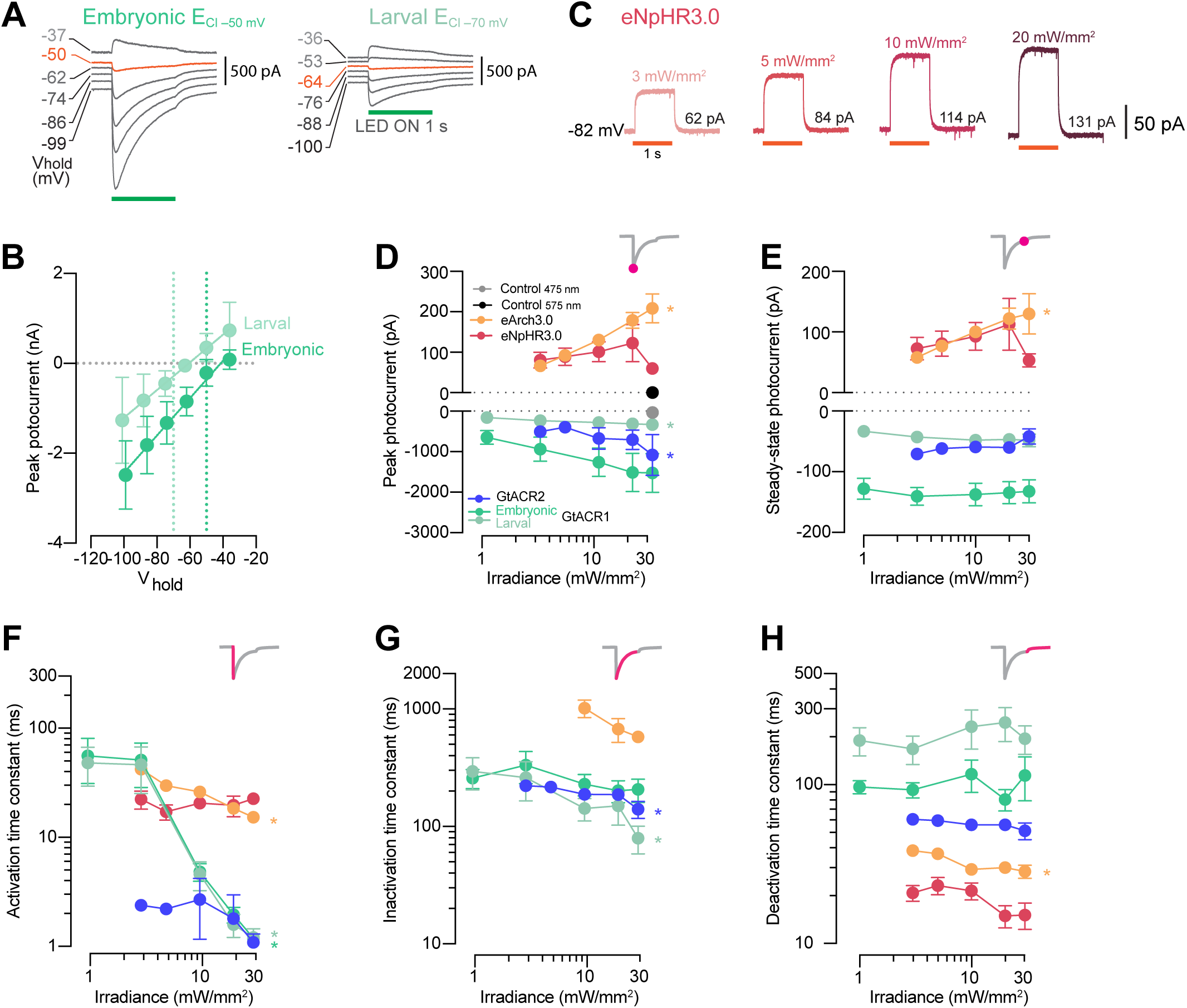
Photocurrent properties vs. irradiance. **A** Example GtACR1 photocurrents obtained by providing a 1 s light periods at different holding potentials (V_hold_) using intracellular solutions approximating either embryonic or larval ECl. Orange traces denote holding potentials closest to ECl. **B** GtACR1 photocurrent I-V curves (mean ± SD). Photocurrents reverse with a positive 5–10 mV shift relative to ECl (dotted lines) in both solutions. **C** Example photocurrents from an eNpHR3.0-expressing cell at different irradiance levels (3– 20 mW/mm^2^). **D,E** Photocurrent peak (**D**) and steady-state (**E**) amplitude vs. irradiance (mean ± SEM, across cells). Asterisks indicate a significant non-zero slope. **F–H** Photocurrent activation (**F**), inactivation (**G**) and deactivation (**H**) time constants vs. irradiance (mean ± SEM). eNpHR3.0 photocurrents did not inactivate hence no inactivation time constant was computed.

**Figure 9-figure supplement 1.**
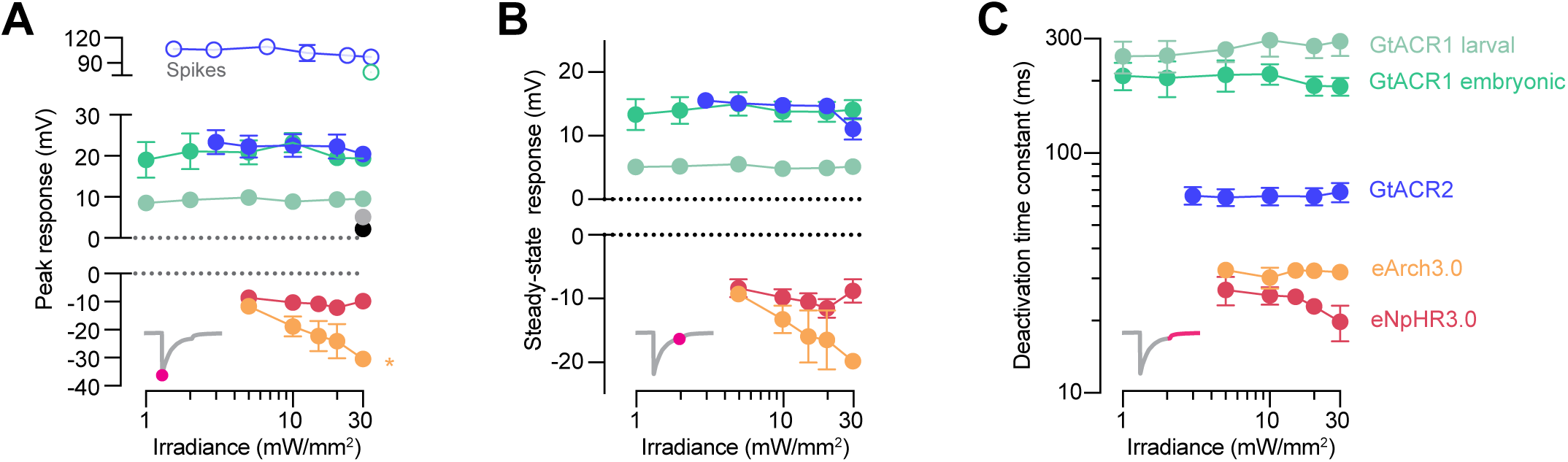
Optogenetically-evoked voltage responses vs. irradiance. ** A–C** Peak (**A**) and steady-state (**B**) responses and deactivation time constant (**C**) of voltage deflections vs. irradiance (mean ± SEM, across cells). eArch3.0 was the only opsin showing irradiance-dependent modulation of peak voltage response.

## References

1. Adamantidis, A., Arber, S., Bains, J.S., Bamberg, E., Bonci, A., Buzsaki, G., Cardin, J.A., Costa, R.M., Dan, Y., Goda, Y., et al. (2015). Optogenetics: 10 years after ChR2 in neurons--views from the community. Nat Neurosci 18, 1202–1212.

2. Andalman, A.S., Burns, V.M., Lovett-Barron, M., Broxton, M., Poole, B., Yang, S.J., Grosenick, L., Lerner, T.N., Chen, R., Benster, T., et al. (2019). Neuronal Dynamics Regulating Brain and Behavioral State Transitions. Cell 177, 970–985.e920.

3. Antinucci, P., Folgueira, M., and Bianco, I.H. (2019). Pretectal neurons control hunting behaviour. Elife 8.

4. Arrenberg, A.B., Del Bene, F., and Baier, H. (2009). Optical control of zebrafish behavior with halorhodopsin. Proc Natl Acad Sci U S A 106, 17968–17973.

5. Arrenberg, A.B., and Driever, W. (2013). Integrating anatomy and function for zebrafish circuit analysis. Front Neural Circuits 7, 74.

6. Asakawa, K., and Kawakami, K. (2008). Targeted gene expression by the Gal4-UAS system in zebrafish. Dev Growth Differ 50, 391–399.

7. Asakawa, K., Suster, M.L., Mizusawa, K., Nagayoshi, S., Kotani, T., Urasaki, A., Kishimoto, Y., Hibi, M., and Kawakami, K. (2008). Genetic dissection of neural circuits by Tol2 transposon-mediated Gal4 gene and enhancer trapping in zebrafish. Proc Natl Acad Sci U S A 105, 1255–1260.

8. Balciunas, D., Davidson, A.E., Sivasubbu, S., Hermanson, S.B., Welle, Z., and Ekker, S.C. (2004). Enhancer trapping in zebrafish using the Sleeping Beauty transposon. BMC Genomics 5, 62.

9. Beattie, C.E., Hatta, K., Halpern, M.E., Liu, H., Eisen, J.S., and Kimmel, C.B. (1997). Temporal separation in the specification of primary and secondary motoneurons in zebrafish. Dev Biol 187, 171–182.

10. Bello-Rojas, S., Istrate, A.E., Kishore, S., and McLean, D.L. (2019). Central and peripheral innervation patterns of defined axial motor units in larval zebrafish. J Comp Neurol 527, 2557–2572.

11. Ben Fredj, N., Hammond, S., Otsuna, H., Chien, C.B., Burrone, J., and Meyer, M.P. (2010). Synaptic activity and activity-dependent competition regulates axon arbor maturation, growth arrest, and territory in the retinotectal projection. J Neurosci 30, 10939–10951.

12. Ben-Ari, Y. (2002). Excitatory actions of gaba during development: the nature of the nurture. Nat Rev Neurosci 3, 728–739.

13. Bernal Sierra, Y.A., Rost, B.R., Pofahl, M., Fernandes, A.M., Kopton, R.A., Moser, S., Holtkamp, D., Masala, N., Beed, P., Tukker, J.J., et al. (2018). Potassium channel-based optogenetic silencing. Nat Commun 9, 4611.

14. Berndt, A., Lee, S.Y., Wietek, J., Ramakrishnan, C., Steinberg, E.E., Rashid, A.J., Kim, H., Park, S., Santoro, A., Frankland, P.W., et al. (2016). Structural foundations of optogenetics: Determinants of channelrhodopsin ion selectivity. Proc Natl Acad Sci U S A 113, 822–829.

15. Berndt, A., Schoenenberger, P., Mattis, J., Tye, K.M., Deisseroth, K., Hegemann, P., and Oertner, T.G. (2011). High-efficiency channelrhodopsins for fast neuronal stimulation at low light levels. Proc Natl Acad Sci U S A 108, 7595–7600.

16. Bohm, U.L., Prendergast, A., Djenoune, L., Nunes Figueiredo, S., Gomez, J., Stokes, C., Kaiser, S., Suster, M., Kawakami, K., Charpentier, M., et al. (2016). CSF-contacting neurons regulate locomotion by relaying mechanical stimuli to spinal circuits. Nat Commun 7, 10866.

17. Boyden, E.S. (2011). A history of optogenetics: the development of tools for controlling brain circuits with light. F1000 Biol Rep 3, 11.

18. Boyden, E.S. (2015). Optogenetics and the future of neuroscience. Nat Neurosci 18, 1200–1201.

19. Boyden, E.S., Zhang, F., Bamberg, E., Nagel, G., and Deisseroth, K. (2005). Millisecond-timescale, genetically targeted optical control of neural activity. Nat Neurosci 8, 1263–1268.

20. Chow, B.Y., Han, X., Dobry, A.S., Qian, X., Chuong, A.S., Li, M., Henninger, M.A., Belfort, G.M., Lin, Y., Monahan, P.E., et al. (2010). High-performance genetically targetable optical neural silencing by light-driven proton pumps. Nature 463, 98–102.

21. Deisseroth, K. (2015). Optogenetics: 10 years of microbial opsins in neuroscience. Nat Neurosci 18, 1213–1225.

22. Deisseroth, K., and Hegemann, P. (2017). The form and function of channelrhodopsin. Science 357.

23. Del Bene, F., and Wyart, C. (2012). Optogenetics: a new enlightenment age for zebrafish neurobiology. Dev Neurobiol 72, 404–414.

24. Douglass, A.D., Kraves, S., Deisseroth, K., Schier, A.F., and Engert, F. (2008). Escape behavior elicited by single, channelrhodopsin-2-evoked spikes in zebrafish somatosensory neurons. Curr Biol 18, 1133–1137.

25. Drapeau, P., Ali, D.W., Buss, R.R., and Saint-Amant, L. (1999). In vivo recording from identifiable neurons of the locomotor network in the developing zebrafish. J Neurosci Methods 88, 1–13.

26. Fidelin, K., Djenoune, L., Stokes, C., Prendergast, A., Gomez, J., Baradel, A., Del Bene, F., and Wyart, C. (2015). State-Dependent Modulation of Locomotion by GABAergic Spinal Sensory Neurons. Curr Biol 25, 3035–3047.

27. Forster, D., Dal Maschio, M., Laurell, E., and Baier, H. (2017). An optogenetic toolbox for unbiased discovery of functionally connected cells in neural circuits. Nat Commun 8, 116.

28. Friedmann, D., Hoagland, A., Berlin, S., and Isacoff, E.Y. (2015). A spinal opsin controls early neural activity and drives a behavioral light response. Curr Biol 25, 69–74.

29. Fujimoto, E., Gaynes, B., Brimley, C.J., Chien, C.B., and Bonkowsky, J.L. (2011). Gal80 intersectional regulation of cell-type specific expression in vertebrates. Dev Dyn 240, 2324–2334.

30. Govorunova, E.G., Sineshchekov, O.A., Janz, R., Liu, X., and Spudich, J.L. (2015). Natural light-gated anion channels: A family of microbial rhodopsins for advanced optogenetics. Science 349, 647–650.

31. Gradinaru, V., Thompson, K.R., Zhang, F., Mogri, M., Kay, K., Schneider, M.B., and Deisseroth, K. (2007). Targeting and readout strategies for fast optical neural control in vitro and in vivo. J Neurosci 27, 14231–14238.

32. Gradinaru, V., Zhang, F., Ramakrishnan, C., Mattis, J., Prakash, R., Diester, I., Goshen, I., Thompson, K.R., and Deisseroth, K. (2010). Molecular and cellular approaches for diversifying and extending optogenetics. Cell 141, 154–165.

33. Hochbaum, D.R., Zhao, Y., Farhi, S.L., Klapoetke, N., Werley, C.A., Kapoor, V., Zou, P., Kralj, J.M., Maclaurin, D., Smedemark-Margulies, N., et al. (2014). All-optical electrophysiology in mammalian neurons using engineered microbial rhodopsins. Nat Methods 11, 825–833.

34. Horstick, E.J., Jordan, D.C., Bergeron, S.A., Tabor, K.M., Serpe, M., Feldman, B., and Burgess, H.A. (2015). Increased functional protein expression using nucleotide sequence features enriched in highly expressed genes in zebrafish. Nucleic Acids Res 43, e48.

35. Huber, D., Petreanu, L., Ghitani, N., Ranade, S., Hromadka, T., Mainen, Z., and Svoboda, K. (2008). Sparse optical microstimulation in barrel cortex drives learned behaviour in freely moving mice. Nature 451, 61–64.

36. Kikuta, H., and Kawakami, K. (2009). Transient and stable transgenesis using tol2 transposon vectors. Methods Mol Biol 546, 69–84.

37. Kimmel, C.B., Hatta, K., and Metcalfe, W.K. (1990). Early axonal contacts during development of an identified dendrite in the brain of the zebrafish. Neuron 4, 535–545.

38. Klapoetke, N.C., Murata, Y., Kim, S.S., Pulver, S.R., Birdsey-Benson, A., Cho, Y.K., Morimoto, T.K., Chuong, A.S., Carpenter, E.J., Tian, Z., et al. (2014). Independent optical excitation of distinct neural populations. Nat Methods 11, 338–346.

39. Li, N., Chen, S., Guo, Z.V., Chen, H., Huo, Y., Inagaki, H.K., Chen, G., Davis, C., Hansel, D., Guo, C., et al. (2019). Spatiotemporal constraints on optogenetic inactivation in cortical circuits. Elife 8.

40. Lister, J.A., Robertson, C.P., Lepage, T., Johnson, S.L., and Raible, D.W. (1999). nacre encodes a zebrafish microphthalmia-related protein that regulates neural-crest-derived pigment cell fate. Development 126, 3757–3767.

41. Luo, L., Callaway, E.M., and Svoboda, K. (2008). Genetic dissection of neural circuits. Neuron 57, 634–660.

42. Mahn, M., Gibor, L., Patil, P., Cohen-Kashi Malina, K., Oring, S., Printz, Y., Levy, R., Lampl, I., and Yizhar, O. (2018). High-efficiency optogenetic silencing with soma-targeted anion-conducting channelrhodopsins. Nat Commun 9, 4125.

43. Mahn, M., Prigge, M., Ron, S., Levy, R., and Yizhar, O. (2016). Biophysical constraints of optogenetic inhibition at presynaptic terminals. Nat Neurosci 19, 554–556.

44. Malyshev, A.Y., Roshchin, M.V., Smirnova, G.R., Dolgikh, D.A., Balaban, P.M., and Ostrovsky, M.A. (2017). Chloride conducting light activated channel GtACR2 can produce both cessation of firing and generation of action potentials in cortical neurons in response to light. Neurosci Lett 640, 76–80.

45. Mardinly, A.R., Oldenburg, I.A., Pegard, N.C., Sridharan, S., Lyall, E.H., Chesnov, K., Brohawn, S.G., Waller, L., and Adesnik, H. (2018). Precise multimodal optical control of neural ensemble activity. Nat Neurosci 21, 881–893.

46. Margrie, T.W., Brecht, M., and Sakmann, B. (2002). In vivo, low-resistance, whole-cell recordings from neurons in the anaesthetized and awake mammalian brain. Pflugers Arch 444, 491–498.

47. Mattis, J., Tye, K.M., Ferenczi, E.A., Ramakrishnan, C., O’Shea, D.J., Prakash, R., Gunaydin, L.A., Hyun, M., Fenno, L.E., Gradinaru, V., et al. (2011). Principles for applying optogenetic tools derived from direct comparative analysis of microbial opsins. Nat Methods 9, 159–172.

48. Menelaou, E., and McLean, D.L. (2012). A gradient in endogenous rhythmicity and oscillatory drive matches recruitment order in an axial motor pool. J Neurosci 32, 10925–10939.

49. Miesenbock, G. (2009). The optogenetic catechism. Science 326, 395–399.

50. Miesenbock, G. (2011). Optogenetic control of cells and circuits. Annu Rev Cell Dev Biol 27, 731–758.

51. Mohamed, G.A., Cheng, R.K., Ho, J., Krishnan, S., Mohammad, F., Claridge-Chang, A., and Jesuthasan, S. (2017). Optical inhibition of larval zebrafish behaviour with anion channelrhodopsins. BMC Biol 15, 103.

52. Portugues, R., Severi, K.E., Wyart, C., and Ahrens, M.B. (2013). Optogenetics in a transparent animal: circuit function in the larval zebrafish. Curr Opin Neurobiol 23, 119–126.

53. Prigge, M., Schneider, F., Tsunoda, S.P., Shilyansky, C., Wietek, J., Deisseroth, K., and Hegemann, P. (2012). Color-tuned channelrhodopsins for multiwavelength optogenetics. J Biol Chem 287, 31804–31812.

54. Reynolds, A., Brustein, E., Liao, M., Mercado, A., Babilonia, E., Mount, D.B., and Drapeau, P. (2008). Neurogenic role of the depolarizing chloride gradient revealed by global overexpression of KCC2 from the onset of development. J Neurosci 28, 1588–1597.

55. Sagasti, A., Guido, M.R., Raible, D.W., and Schier, A.F. (2005). Repulsive interactions shape the morphologies and functional arrangement of zebrafish peripheral sensory arbors. Curr Biol 15, 804–814.

56. Saint-Amant, L., and Drapeau, P. (1998). Time course of the development of motor behaviors in the zebrafish embryo. J Neurobiol 37, 622–632.

57. Saint-Amant, L., and Drapeau, P. (2000). Motoneuron activity patterns related to the earliest behavior of the zebrafish embryo. J Neurosci 20, 3964–3972.

58. Saint-Amant, L., and Drapeau, P. (2003). Whole-cell patch-clamp recordings from identified spinal neurons in the zebrafish embryo. Methods Cell Sci 25, 59–64.

59. Scheer, N., and Campos-Ortega, J.A. (1999). Use of the Gal4-UAS technique for targeted gene expression in the zebrafish. Mech Dev 80, 153–158.

60. Schneider, F., Grimm, C., and Hegemann, P. (2015). Biophysics of Channelrhodopsin. Annu Rev Biophys 44, 167–186.

61. Scott, E.K., Mason, L., Arrenberg, A.B., Ziv, L., Gosse, N.J., Xiao, T., Chi, N.C., Asakawa, K., Kawakami, K., and Baier, H. (2007). Targeting neural circuitry in zebrafish using GAL4 enhancer trapping. Nat Methods 4, 323–326.

62. Sjulson, L., Cassataro, D., DasGupta, S., and Miesenbock, G. (2016). Cell-Specific Targeting of Genetically Encoded Tools for Neuroscience. Annu Rev Genet 50, 571–594.

63. Song, J., Ampatzis, K., Bjornfors, E.R., and El Manira, A. (2016). Motor neurons control locomotor circuit function retrogradely via gap junctions. Nature 529, 399–402.

64. Suster, M.L., Abe, G., Schouw, A., and Kawakami, K. (2011). Transposon-mediated BAC transgenesis in zebrafish. Nat Protoc 6, 1998–2021.

65. Warp, E., Agarwal, G., Wyart, C., Friedmann, D., Oldfield, C.S., Conner, A., Del Bene, F., Arrenberg, A.B., Baier, H., and Isacoff, E.Y. (2012). Emergence of patterned activity in the developing zebrafish spinal cord. Curr Biol 22, 93–102.

66. Williams, R.H., Tsunematsu, T., Thomas, A.M., Bogyo, K., Yamanaka, A., and Kilduff, T.S. (2019). Transgenic Archaerhodopsin-3 Expression in Hypocretin/Orexin Neurons Engenders Cellular Dysfunction and Features of Type 2 Narcolepsy. J Neurosci.

67. Wyart, C., Del Bene, F., Warp, E., Scott, E.K., Trauner, D., Baier, H., and Isacoff, E.Y. (2009). Optogenetic dissection of a behavioural module in the vertebrate spinal cord. Nature 461, 407–410.

68. Yizhar, O., Fenno, L.E., Davidson, T.J., Mogri, M., and Deisseroth, K. (2011). Optogenetics in neural systems. Neuron 71, 9–34.

69. Zhang, F., Wang, L.P., Brauner, M., Liewald, J.F., Kay, K., Watzke, N., Wood, P.G., Bamberg, E., Nagel, G., Gottschalk, A., et al. (2007). Multimodal fast optical interrogation of neural circuitry. Nature 446, 633–639.

70. Zhang, R.W., Wei, H.P., Xia, Y.M., and Du, J.L. (2010). Development of light response and GABAergic excitation-to-inhibition switch in zebrafish retinal ganglion cells. J Physiol 588, 2557–2569.

